# Kir4.2 deficiency drives progressive Parkinson’s disease-like motor, cognitive and neuropathological phenotypes in mice

**DOI:** 10.64898/2026.03.01.708908

**Authors:** Benjamin Garland, Zhiqiang Shen, Mo Chen, Bingmiao Gao, Kailin Mao, Des R. Richardson, George D. Mellick, Linlin Ma

**Affiliations:** School of Environment and Science, Griffith University, Brisbane 4111, Queensland, Australia; Institute for Biomedicine and Glycomics, Griffith University, Brisbane 4111, Queensland, Australia; Engineering Research Center of Tropical Medicine Innovation and Transformation of Ministry of Education, Hainan Key Laboratory for Research and Development of Tropical Herbs, College of Pharmacy, Hainan Medical University, Haikou 571199, Hainan, China; Center for Cancer Cell Biology and Drug Discovery, Institute for Biomedicine and Glycomics, Griffith University, Gold Coast 4215, Queensland, Australia

**Author notes:** B. Garland & Z. Shen are equal first authors. **Authors for correspondence: Linlin Ma, MD & PhD**, School of Environment and Science, Institute for Biomedicine and Glycomics, Griffith University, Brisbane, Queensland, Australia, George D. Mellick, PhD, School of Environment and Science, Institute for Biomedicine and Glycomics, Griffith University, Brisbane, Queensland, Australia, **Des R. Richardson**, **PhD.**, **D.Sc**., Center for Cancer Cell Biology and Drug Discovery, Institute for Biomedicine and Glycomics, Griffith University, Gold Coast, Queensland, Australia.

**Keywords:** Parkinson’s disease, inwardly rectifying potassium channel, Kir4.2, *KCNJ15*, motor syndrome, anxiety, cognitive decline, neuroinflammation, synucleinopathy

## Abstract

A critical barrier to therapeutic development in Parkinson’s disease (PD) is the lack of endogenous genetic models that spontaneously recapitulate the disease’s progressive, anatomically selective nigrostriatal pathology. Genetic studies have linked *KCNJ15*, encoding the inwardly rectifying potassium channel Kir4.2, to familial PD via a loss-of-function dominant-negative variant; however, its mechanistic role in neurodegeneration remains unexplored. Here, we demonstrate that *Kcnj15⁻/⁻* mice spontaneously develop a progressive PD-like phenotype, exhibiting a “coordination-first” motor syndrome, anxiety-like behavioural changes and spatial memory impairments. Crucially, neuropathology reveals selective degeneration of substantia nigra pars compacta neurons, sparing the ventral tegmental area, with marked microglial hyperactivation and phosphorylated α-synuclein accumulation, faithfully recapitulating the topography of human PD. Striatal transcriptomics further reveals upregulation of oligodendrocyte- and myelin-associated genes, implicating compensatory glial remodeling. These findings identify Kir4.2 as a critical homeostatic regulator of nigrostriatal integrity and establish the *Kcnj15⁻/⁻* mouse as a new physiologically accurate and translatable model for PD research.

## Introduction

Parkinson’s disease (PD) has become the fastest-growing neurodegenerative disorder and a leading cause of chronic disability, with its prevalence projected to more than double by 2050 [1]. Although clinically defined by cardinal motor features, such as bradykinesia, tremor, and rigidity, PD is increasingly recognized as a systemic circuitopathy. Within this framework, a constellation of non-motor symptoms (e.g., cognitive decline, anxiety, sleep disruption, and autonomic dysfunction) frequently precedes motor onset and contributes substantially to overall disease burden [2]. Because the molecular mechanisms underlying this broad phenotypic spectrum remain elusive, identifying novel molecular drivers that initiate or accelerate the neurodegenerative cascade is essential for understanding and ultimately modifying this debilitating disease.

A critical barrier to therapeutic progress in PD is the lack of preclinical animal models that faithfully recapitulate its progressive etiology. Widely used neurotoxin-based models, such as 6-OHDA and MPTP models, rely on acute pharmacological destruction of the nigrostriatal pathway, thereby bypassing the upstream genetic and molecular events that initiate the disease, limiting their utility for studying prodromal or early-stage pathology [2, 3]. Conversely, transgenic models, such as transgenic mice overexpressing mutant human α-synuclein or knockouts of autosomal recessive PD genes, often fail to exhibit the spontaneous, anatomically selective dopaminergic neurodegeneration in the substantia nigra pars compacta (SNpc) that defines human PD [3–7]. The absence of an endogenous genetic model that genuinely mimics the human disease process fundamentally constrains our ability to study disease initiation, identify early intervention windows, and test neurorestorative therapies.

In previous genetic linkage studies of a four-generation family with familial PD, we identified *KCNJ15* as a candidate gene that strongly segregated with disease status [8]. Subsequent *in vitro* functional characterization demonstrated that the identified mutation acts as a loss-of-function variant with dominant-negative effects on its encoded protein, the inwardly rectifying potassium channel Kir4.2 [9].

Inwardly rectifying potassium (Kir) channels are fundamental regulators of cellular excitability, essential for maintaining K^+^ homeostasis, setting the resting membrane potential, and shaping action potential duration [10, 11]. Dysfunction of Kir channels is increasingly implicated in neurodegenerative disorders, including Alzheimer’s, Huntington’s, and PD [12, 13]. However, very little is known about Kir4.2, particularly in the nervous system.

Given the direct genetic link between Kir4.2 loss-of-function and familial PD [8], we hypothesized that Kir4.2 is not merely a passive genetic marker, but an active regulator of nigrostriatal integrity. To test this, we generated *Kcnj15*^⁻/⁻^ mice and performed longitudinal behavioural, neuropathological, and transcriptomic profiling across an aging trajectory of 6 to 18 months. We report that loss of Kir4.2 drives a comprehensive PD-like phenotype, encompassing progressive motor coordination deficits, spatial memory impairment, and dynamic anxiety-like behaviors. Crucially, these behavioral deficits are accompanied by anatomically selective pathology characterized by nigral microglial hyperactivation, pathological α-synuclein accumulation, and dopaminergic degeneration. Furthermore, transcriptomic profiling further revealed a distinct striatal signature of oligodendrocyte and myelin remodeling, suggesting a complex neuroglial response to channel loss. Collectively, our findings establish the *Kcnj15*^⁻/⁻^ mouse as a translatable genetic model of spontaneous, topographically faithful nigrostriatal neurodegeneration. These data identify Kir4.2 as a new homeostatic regulator of nigral health and provide a mechanistic foundation for exploring Kir4.2-dependent neuroimmune and myelin-support pathways as therapeutic targets in PD.

## Results

### *Kcnj15⁻/⁻* mice exhibit progressive coordination-first motor deficits

Aging is the strongest risk factor for PD, with motor symptoms typically emerging progressively in both patients and rodent models [14]. To determine whether Kir4.2 deficiency recapitulates this trajectory, we performed a longitudinal behavioral assessment in *Kcnj15*^⁻/⁻^ mice at both six and twelve months of age (**Fig. 1**, *n* = 9-22). These time points were selected to model early and more advanced stages of phenotypic progression in rodent models [15].

**Figure 1.**
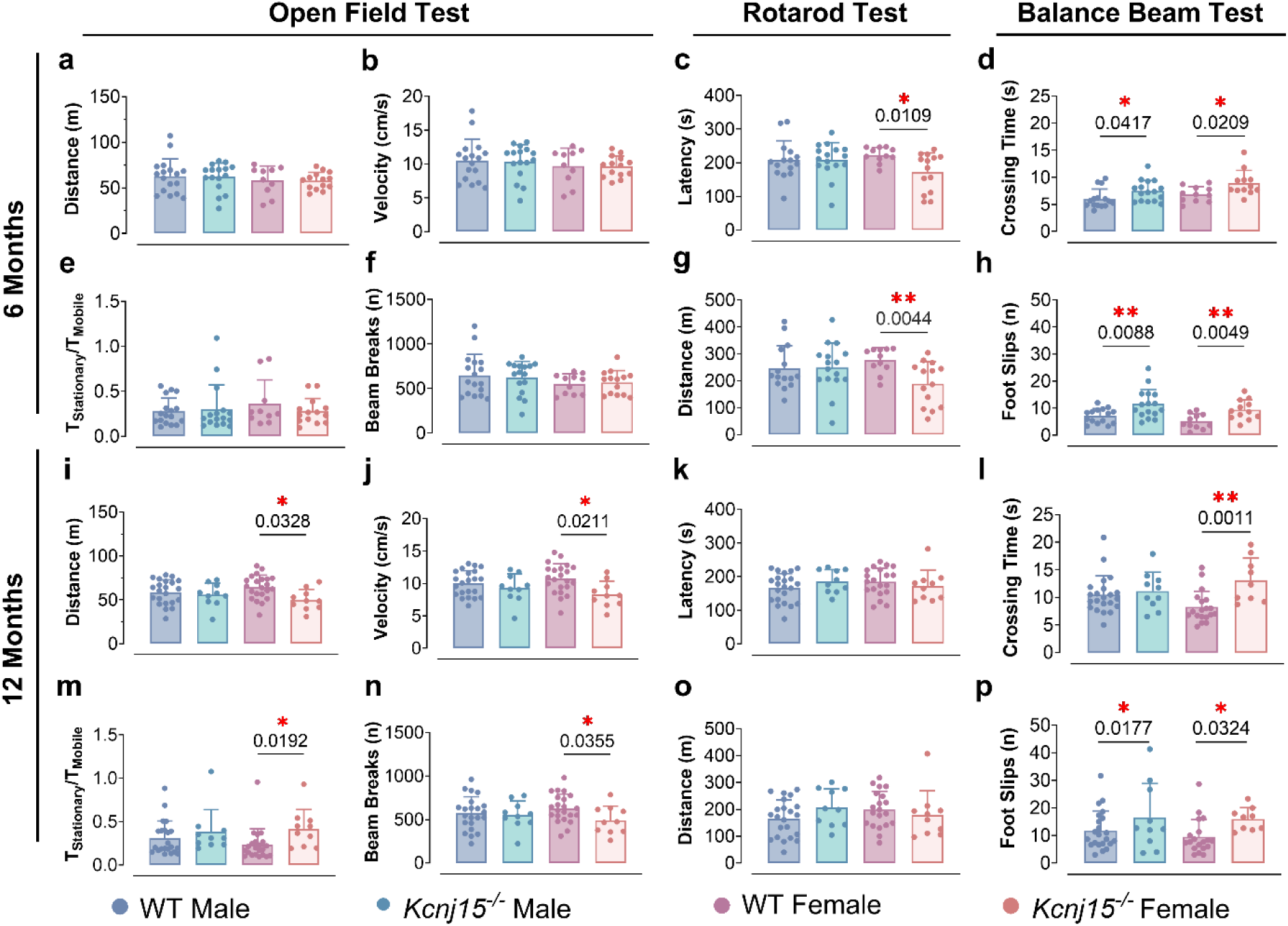
Behavioral assessments reveal age-dependent motor function deficits and reduced activity in *Kcnj15^⁻/⁻^* mice. A battery of behavioral tests was conducted on wild-type (WT) and *Kcnj15* knockout (*Kcnj15*^-/-^) mice at six months of age (**a-h**) and twelve months of age (**i-p**). **Open field test**: Total distance travelled (**a & i**); average velocity during the mobile phase (**b & j**); the ratio of stationary to mobile time (**e & m**); and beam-break frequency (**f & n**) were measured to evaluate locomotor activity. **Accelerating rotarod test**: Mice were trained to run on a horizontal rotarod rotating at an accelerating speed from 4 to 60 rpm over 300 s, after which the speed was maintained at 60 rpm until the mouse fell. Latency to fall (**c & k**) and total travel distance (**g & o**) were recorded to assess motor coordination, balance, endurance, and performance capacity. **Balance beam test**: Time taken to traverse a 50 mm-thick beam (**d & l**) and the number of foot slips (**h & p**) were quantified to assess coordination and balance. Each dot represents an individual mouse; error bars show mean ± SD. Statistical comparisons were performed using unpaired two-tailed t-tests. \**P* < 0.05, \*\**P* < 0.01. At six months of age, WT male: *n =* 15-17, *Kcnj15*^⁻/⁻^ male: *n =* 17, WT female: *n =* 11, *Kcnj15*^⁻/⁻^ female: *n =* 12-14; at twelve months of age, WT male: *n =* 22, *Kcnj15*^⁻/⁻^ male: *n =* 10, WT female: *n =* 19, *Kcnj15*^⁻/⁻^ female: *n =* 9-10.

Given that PD is clinically defined by bradykinesia, rigidity, gait disturbances, and postural instability, we first examined general locomotor activity using the open-field test in a low-stress, unconfined setting over a prolonged 10 min period. At 6 months of age, *Kcnj15*^⁻/⁻^ mice were indistinguishable from WT controls across all metrics, including mean distance travelled (**Fig. 1a**), average velocity during the mobile phase (**Fig. 1b**), the ratio of stationary to mobile time (**Fig. 1e**), and beam-break frequency (**Fig. 1f**). These data indicate that Kir4.2 is not required for basal locomotor output in early adulthood. By contrast, by 12 months of age, female *Kcnj15*^⁻/⁻^ mice displayed a significant decline in spontaneous activity, evidenced by decreased distance travelled (*P* = 0.0328; **Fig. 1i**), lower velocity (*P* = 0.0211; **Fig. 1j**), increased immobility (*P* = 0.0192; **Fig. 1m**), and fewer beam breaks (*P* = 0.0355; **Fig. 1n**). No comparable open-field deficits were evident in 12-month-old males. These data indicate that broad locomotor impairment emerges later and is most prominent in females. This age-dependent deterioration is consistent with the progressive nature of PD-related motor dysfunction [2].

To probe the integrity of the nigrostriatal and cerebellar circuits required for complex motor coordination, we employed the accelerating rotarod test. This is a standard assay used to uncover cerebellar, basal ganglia, and motor-cortical impairments in models of neurodegenerative diseases, ataxia, and motor impairment [16]. At 6 months old, female *Kcnj15*^⁻/⁻^ mice displayed significant motor impairments, with shorter latency to fall (*P* = 0.0109; **Fig. 1c**) and reduced cumulative distance travelled across trials (*P* = 0.0044; **Fig. 1g**), indicating early deficits in coordination and motor endurance that precede the later decline seen in the open field. Notably, this deficit occurs despite a significant drop in their overall body weight compared to WT controls (*P* = 0.0006, **Supplementary Fig. 2**). Because increased body weight generally places a biomechanical disadvantage on rotarod performance [17], the lighter mass of the KO females should theoretically serve as an advantage. The fact that *Kcnj15*^⁻/⁻^ females perform significantly worse despite this physical advantage strongly suggests a profound, genotype-driven neurological deficit that overpowers their reduced body weight.

While this phenotype appeared to resolve at 12 months (**Fig. 1k & 1o**), rotarod performance is inherently confounded by body weight dynamics. As shown in **Supplementary Figure 2**, both male and female *Kcnj15*^⁻/⁻^ mice were significantly smaller than WT controls at both 6 and 12 months (*P* = 0.0006-0.008). To explicitly decouple the effects of this reduced body size from true motor deficits, we performed multiple linear regression analyses incorporating weight as a covariate (**Fig. 2**). In male mice, the analysis results revealed a significant interaction between genotype and body weight (Weight × Genotype Interaction *P* = 0.0307; **Fig. 2a**). While WT males exhibited a significant negative correlation between body weight and rotarod performance (where heavier mice fall faster, R^2^ = 0.5992), *Kcnj15* deletion completely uncoupled this relationship (R^2^ = 0.0037; **Fig. 2a**). This suggests that *Kcnj15*^⁻/⁻^ modifies weight-dependent rotarod vulnerability in males, and this trend persisted at 12 months (Interaction *P* = 0.0646; **Fig. 2c**). Because lightweight WT and *Kcnj15*^⁻/⁻^ males perform similarly well, the true functional divergence between the genotypes is obscured in simple group averages and only becomes apparent as the mice face the biomechanical stress of increased body weight.

**Figure 2.**
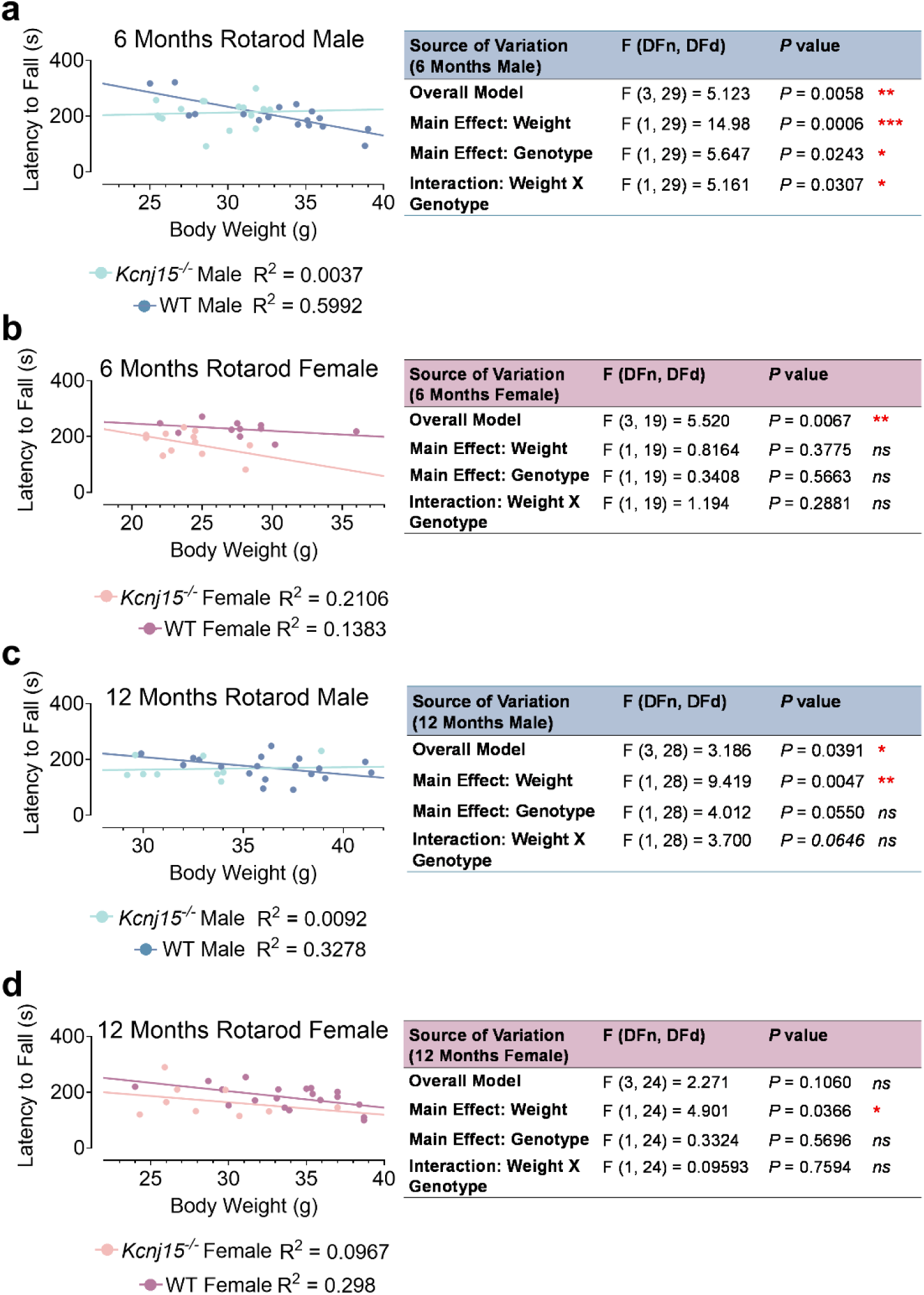
*Kcnj15* deletion alters the relationship between body weight and rotarod performance in male mice. **(a–d)** Scatter plots illustrating the relationship between body weight (g) and rotarod performance (Latency to Fall, s) across age and sex cohorts: 6-month-old males **(a)**, 6-month-old females **(b)**, 12-month-old males **(c)**, and 12-month-old females **(d)**. Individual data points represent single mice for WT (dark circles) and *Kcnj15*^⁻/⁻^ (light circles) groups. Solid lines represent simple linear regression best-fit curves, with corresponding *R²* values shown below each graph. Adjacent tables summarize the results of multiple linear regression analyses evaluating the overall model, the main effects of body weight and genotype, and the weight × genotype interaction for each cohort. *ns*, not significant; *, *P* < 0.05; **, *P* < 0.01; ***, *P* < 0.001.

Conversely, in females, there was no significant interaction between body weight and genotype at either 6 or 12 months (Interaction *P* = 0.2881 and *P* = 0.7594, respectively; **Fig. 2b & 2d**). Because 6-month-old *Kcnj15*^⁻/⁻^ females are significantly lighter than WT controls, this strong collinearity between weight and genotype limits the regression model’s ability to perfectly isolate the two variables (Genotype main effect *P* = 0.5663; **Fig. 2b**). Therefore, while the rotarod highlights a striking vulnerability in *Kcnj15*^⁻/⁻^ females, it cannot definitively uncouple their neurological impairment from their severely altered body weight.

To resolve this, we next examined fine balance, motor coordination, and agility using a more sensitive assay: the elevated balance beam test (**Supplementary Videos 1 & 2**). PD patients characteristically experience declining balance, impaired postural reflexes, and difficulty with coordinated movements, which are symptoms linked to basal ganglia dysfunction and cerebellar-striatal network disruption [18]. Unlike the rotarod, motor performance on the balance beam is not heavily confounded by body weight [19]. Indeed, our multiple linear regression analyses confirmed that body weight was not a significant predictor of latency to cross or foot slips in most cohorts (**Supplementary Fig. 3 and 4**).

At six months of age, both male and female *Kcnj15*^⁻/⁻^ mice took significantly longer to traverse the beam than their WT counterparts (*P* = 0.0417 and *P* = 0.0209, respectively; **Fig. 1d**) and committed a higher frequency of foot slips (*P* = 0.0088 for males, *P* = 0.0049 for females; **Fig. 1h**), demonstrating clear early-onset impairments in balance and coordination. These impairments persisted at 12 months, particularly in females, which continued to exhibit significantly prolonged crossing times (*P* = 0.0011; **Fig. 1l**), and elevated foot slips (*P* = 0.0324; **Fig. 1p**). In 12-month-old males, although overall crossing time had normalized (**Fig. 1l**), foot slip frequency remained significantly higher than in WT controls ( *P* = 0.0177; **Fig. 1p**). Since the balance beam bypasses the body weight limitations of the rotarod, these combined findings confirm that the motor deficits observed in *Kcnj15*^⁻/⁻^ mice across both sexes are largely independent of body weight and are most consistent with genotype-driven impairments in fine motor coordination.

Taken together, these findings show that *Kcnj15* deficiency produces a coordination-first motor phenotype. Deficits in skilled balance and rotarod performance are already evident at 6 months, whereas broader reductions in spontaneous locomotion emerge later, most clearly in 12-month-old females. Thus, loss of Kir4.2 leads to an early and progressive motor syndrome dominated initially by impaired coordination and postural control.

### Spatiotemporal gait parameters are preserved in *Kcnj15*^⁻/⁻^ mice despite coordination deficits

Gait abnormality is another common motor feature of PD, typically characterized by short, narrow steps and a stooped posture, commonly referred to as a “shuffling gait” [2, 20]. To determine whether loss of Kir4.2 affects gait patterning in addition to balance and coordination, we employed the DigiGait™ system (Mouse Specifics) to quantify temporal gait characteristics of each limb and overall gait symmetry in 12-month-old male WT and *Kcnj15*^⁻/⁻^ mice. DigiGait quantifies spatiotemporal gait parameters, including stride length, stances/swing durations, paw placement symmetry, and stride variability, during forced treadmill locomotion [21]. Analysis of temporal parameters across four paws (**Fig. 3a–d, f**) and global gait symmetry (**Fig. 3e**) did not reveal any statistically significant differences between WT and *Kcnj15*^⁻/⁻^ mice. Comprehensive evaluation of 27 DigiGait variables (**Supplementary Figures 5–7**) similarly showed no overt abnormalities.

**Figure 3.**
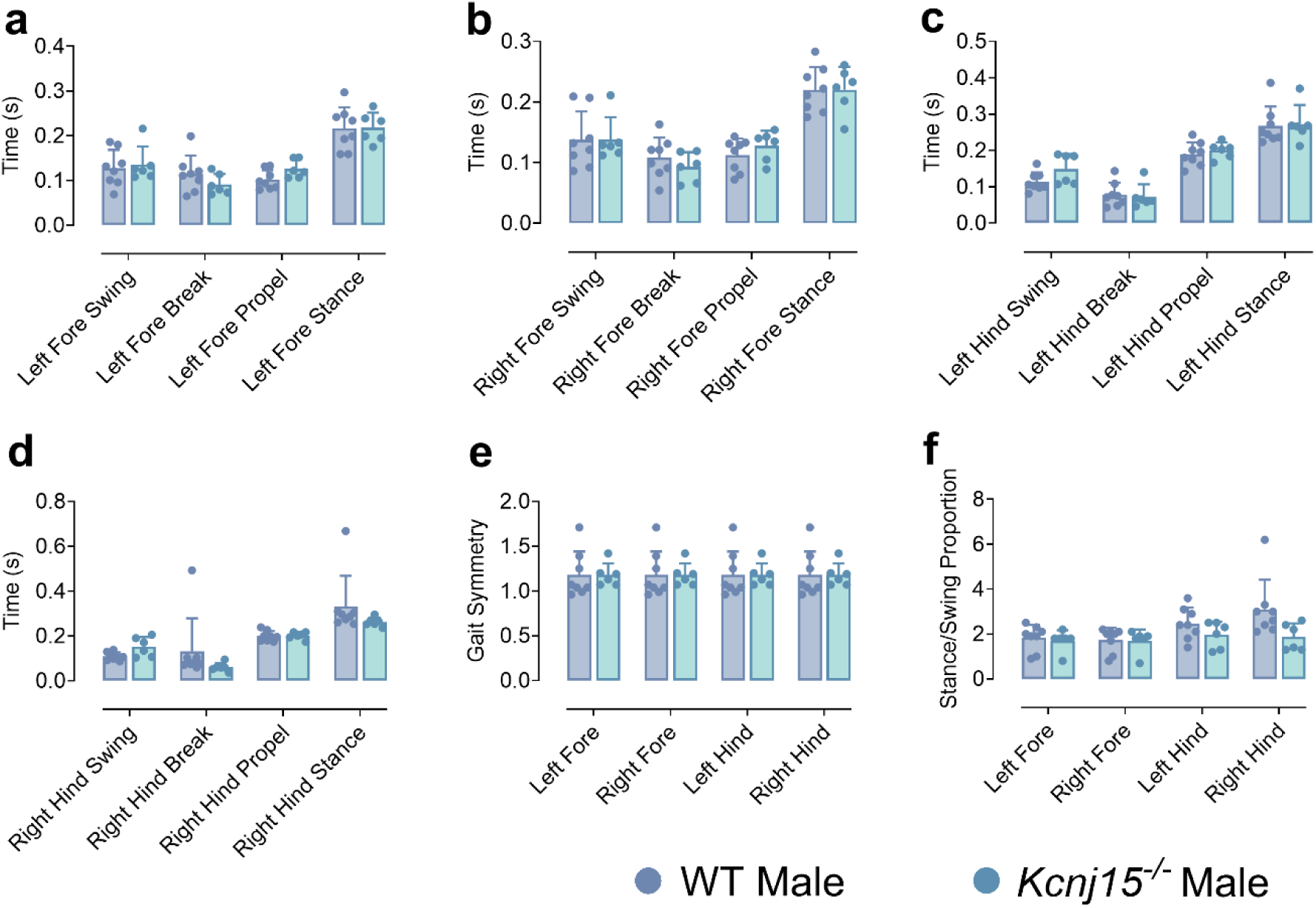
DigiGait analysis shows largely preserved spatiotemporal gait parameters in *Kcnj15*^⁻/⁻^ mice. Automated gait analysis was performed using the DigiGait treadmill system to evaluate locomotor coordination and spatiotemporal gait parameters in wild-type (WT) and *Kcnj15* knockout (*Kcnj15^-/-^*) mice. Temporal gait parameters for all four paws were measured using the DigiGait system in 12-month-old male mice. Bar plots show swing, brake, propel, and stance times for the left fore **(a)**, right fore **(b)**, left hind **(c)**, and right hind **(d)** limbs. The gait symmetry index **(e)** and stance/swing proportion **(f)** are shown for each limb. Each dot represents one mouse. Data were analyzed by unpaired two-tailed *t*-tests and are presented as mean ± SD. WT male: *n =* 8, *Kcnj15*^⁻/⁻^ male: *n =* 6.

DigiGait parameters are typically altered in the presence of spinal or brainstem locomotor circuit dysfunction, significant muscle weakness, or more advanced or asymmetric nigrostriatal degeneration [21, 22]. The largely preserved gait metrics in *Kcnj15*^⁻/⁻^ mice support the view that Kir4.2 loss preferentially impacts higher-order motor integration centers (*e.g.*, basal ganglia-cerebellar loops) before affecting the spinal/brainstem generators of locomotion. This behavioral profile is also consistent with PD progression, as PD-related motor decline typically first manifests in tasks requiring dynamic balance, coordination, and motor planning, while straight, comfortable-pace gait measures may remain relatively preserved [20, 23].

### Loss of Kir4.2 leads to age-progressive long-term spatial memory deficits

Cognitive impairment, particularly in domains of executive function and spatial memory, is now recognized as a core non-motor manifestation of PD that can emerge years before or alongside motor symptoms [2]. The Barnes maze is a well-validated behavioral assay for assessing hippocampal-dependent spatial working memory, special reference memory, and cognitive flexibility in rodents. Compared with more aversive paradigms such as the Morris water maze, the Barnes maze minimises stress and swimming-related confounders, thereby increasing sensitivity to subtle cognitive impairments and reducing the impact of motor deficits on performance [24].

In this task, mice learned to navigate a circular platform with 40 evenly spaced holes along the perimeter (**Fig. 4a**) to locate a single escape box hidden beneath one of the holes within 3 min over a four-day training regime. During this acquisition phase (Days 1–4), *Kcnj15*^⁻/⁻^ mice learned the task effectively, as indicated by the intense red color in the target hole on the heatmap, reflecting longer dwell time (**Fig. 4b**). Acquisition was followed by probe trials on days 5 and 12, in which the escape box was removed, allowing assessment of short - and long-term memory retention, respectively. Rather than focusing on latency and path length, which can be strongly influenced by motivation and locomotor capacity [25], we quantified the distribution of time spent in spatial zones to strengthen interpretation ( **Fig. 4a**). A peaked distribution, with time concentrated in the target and adjacent zones, was interpreted as a memory-guided navigation strategy, whereas a flatter, more uniform distribution reflected random exploration.

**Figure 4.**
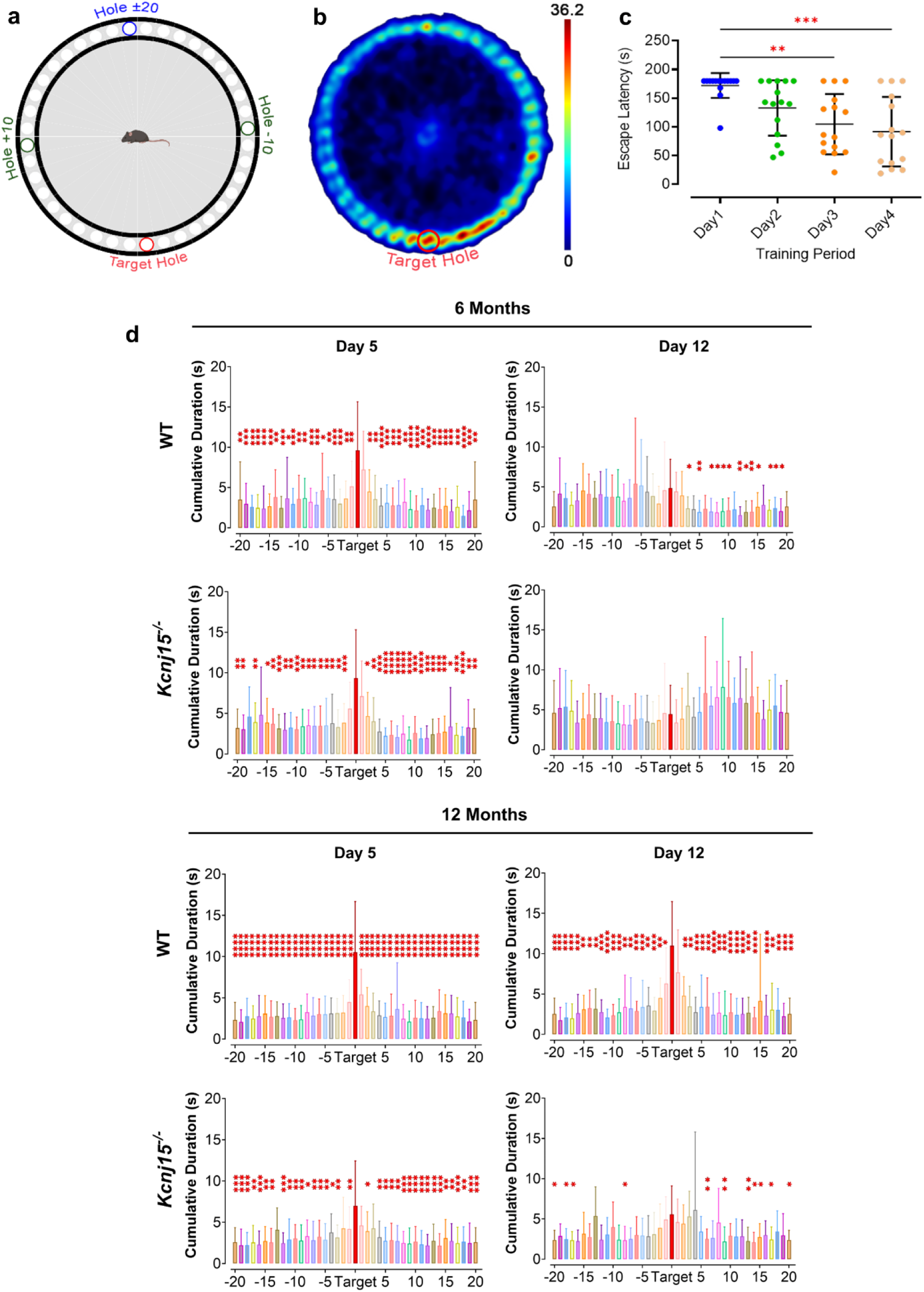
Barnes maze reveals age-dependent long-term spatial memory impairments in *Kcnj15*^⁻/⁻^ mice. **(a)** Schematic of the Barnes maze showing the target/escape hole (red) and the hole-numbering scheme relative to the target. **(b)** Representative spatial occupancy heatmap during the Barnes maze session at 6 months; warmer colours indicate longer dwell time (scale bar). The target hole is marked in red. **(c)** Escape latency across training days (Day 1–Day 4) at 6 months, showing progressive improvement across the training period; each dot represents one animal, and bars indicate mean ± SD. Significance denotes comparisons across days analyzed using one-way ANOVA. (**d & e)** Zone occupancy during probe trials on days 5 and 12 for 6-month-old **(d)** and 12-month-old **(e)** WT and *Kcnj15*^⁻/⁻^ mice. Values are expressed as mean ± SD. The red bar indicates the accumulated dwell time in the target hole. Asterisks denote zones significantly different from the target (repeated measures ANOVA with post hoc comparisons, \*\**P* < 0.01, \*\*\**P* < 0.001, **** *P* < 0.0001). Male and female data were pooled because no significant differences were observed between the male and female cohorts in these tests. At six months of age, WT: *n* = 28, *Kcnj15*^⁻/⁻^: *n* = 31; at twelve months of age, WT: *n* = 41, *Kcnj15*^⁻/⁻^: *n* = 20.

At six months of age, both WT and *Kcnj15*^⁻/⁻^ mice showed robust short-term spatial memory on day 5, spending significantly (*P* < 0.01 to *P* < 0.001) more time in the target zone than in most error zones (**Fig. 4d**). This indicates that initial acquisition and short-term retention of the escape location are preserved in the absence of Kir4.2. By day 12, WT mice exhibited partial decay of spatial memory, as evidenced by fewer error zones differing significantly from the target zone, consistent with normal forgetting over time (**Fig. 4d**). In contrast, *Kcnj15*^⁻/⁻^ mice displayed a largely flat, random pattern of zone exploration with no significant preference for the target location (**Fig. 4d**), indicative of a selective deficit in long-term spatial memory retention rather than a global learning problem.

To investigate whether these deficits are exacerbated by aging, we repeated the test in 12 - month-old mice (**Fig. 4e**). On day 5, both WT and *Kcnj15*^⁻/⁻^ mice again demonstrated good memory of the target location, confirming that even at 12 months, Kir4.2 loss does not prevent acquisition or short-term recall despite their motor impairments (**Fig. 1**). However, by day 12, WT mice maintained a clear preference for the target zone and adjacent zones. In contrast, *Kcnj15*^⁻/⁻^ mice exhibited a more dispersed search pattern with significantly reduced discrimination between the target and error zones (**Fig. 4e**). Although some residual bias towards the former escape location remained, the overall zone occupancy profile of *Kcnj15*^⁻/⁻^ mice was significantly flatter than that of WT controls, indicating a failure to consolidate or retrieve remote spatial memories. Importantly, because both genotypes showed comparable performance at day 5 and general locomotor capacity on the maze did not significantly differ in a way that could explain the selective day-12 deficit, these findings are unlikely to be secondary to gross motor dysfunction or altered motivation.

Taken together, these data show that Kir4.2 deficiency selectively compromises long-term consolidation and/or retrieval of spatial memory, while sparing initial learning or short -term recall, and that this deficit is evident already at 6 months and becomes more pronounced with age. This pattern is consistent with a report that remote spatial memory in the Barnes maze depends on extended hippocampal-cortical networks, which are particularly vulnerable to aging and neurodegenerative processes [26]. Our findings therefore suggest that loss of Kir4.2 shifts the long-term memory circuits towards an early PD-like cognitive vulnerability and accelerates age-related spatial memory decline.

### Age-dependent shifts in anxiety and exploratory strategy in *Kcnj15⁻/⁻* mice

Anxiety is a prevalent prodromal feature of PD, associated with dysregulation of dopaminergic and limbic circuits [27]. We utilized the open field test to dissect anxiety-like behaviors, such as center avoidance [28]. However, anxious mice can also display hyperlocomotion and more center visits, reflecting a stress-induced escape drive rather than true exploratory behavior [29, 30]. Grooming behavior, which represents an additional index of arousal and stress coping, is likewise bidirectional, influenced by opposing mechanisms [31]. On one hand, it serves as a de-arousal/self-soothing behavior when the animal feels sufficiently secure to engage in a vulnerable activity. On the other hand, premature or excessive grooming can signify elevated anxiety or a compulsive-like response [31]. As such, we used a combination of: **(1)** the latency to first center entry; **(2)** time spent in the center zone; and **(3)** latency and frequency of grooming and rearing to dissect anxiety- and arousal-related phenotypes.

At six months of age, *Kcnj15*^⁻/⁻^ males exhibited a significantly increased latency to enter the center of the arena compared to WT controls (*P* = 0.0292; **Fig. 5a**). These results suggested heightened anxiety-like behavior upon exposure to a novel environment [28]. Despite this delayed first center entry, total time spent in the center did not differ between genotypes or sexes (**Fig. 5b**), indicating that once the initial barrier to center exploration was overcome, overall exploratory patterns converged. Additionally, *Kcnj15*^⁻/⁻^ males initiated grooming significantly earlier than both WT males (*P* = 0.0187) and *Kcnj15*^⁻/⁻^ females (*P* = 0.0004; **Fig. 5c**). This shortened grooming latency, in the absence of differences in total grooming or rearing frequency (**Fig. 5d**), suggests altered arousal dynamics rather than simple hyper- or hypo-activity [31, 32].

**Figure 5.**
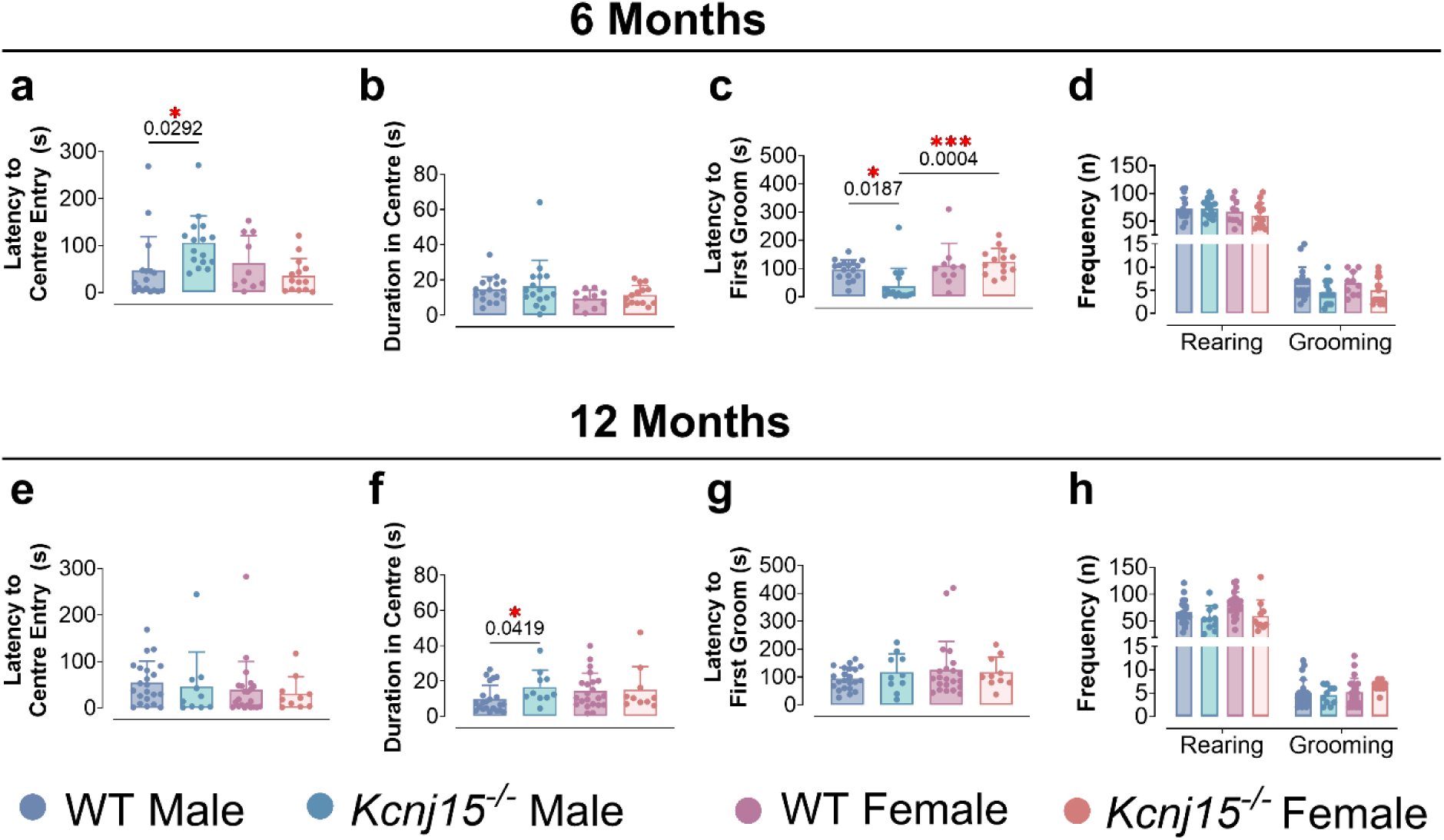
Age- and sex-specific alterations in anxiety-like behaviors in Kir4.2-deficient mice. Anxiety-like behavior and exploratory activity of WT and *Kcnj15*^⁻/⁻^ mice were assessed using the open field test. **(a–d)**, Open field behavior at 6 months. *Kcnj15*^⁻/⁻^ males displayed significantly increased latency to enter the center of the arena compared to WT males (*P* = 0.0292; (**a)**, indicative of heightened anxiety-like behavior. No significant differences were observed in total time spent in the center across genotypes or sexes (**b**). *Kcnj15*^⁻/⁻^ males initiated grooming significantly earlier than WT males (*P* = 0.0187) and *Kcnj15*^⁻/⁻^ females (*P* = 0.0004; **c**). The frequency of grooming and rearing behaviors was not significantly different between groups (**d**). (**e–h)**, Behavioral outcomes at 12 months. Latency to center entry was similar across groups (**e**). *Kcnj15*^⁻/⁻^ males spent significantly more time in the center compared to WT males (*P* = 0.0419; **f**). No differences were detected in grooming latency (**g**) or the frequency of grooming and rearing behaviors (**h**). Data are presented as mean ± SD with each dot representing one mouse. Statistical analysis was performed using one-way ANOVA followed by post hoc tests. These results suggest that *Kcnj15*^⁻/⁻^ males exhibit early anxiety-like phenotypes that shift with age. No significant behavioral alterations were observed in *Kcnj15*^⁻/⁻^ females, indicating a sex-specific effect. At six months of age, WT male: *n* = 17, *Kcnj15*^⁻/⁻^ male: *n* = 16, WT female: *n* = 10, *Kcnj15*^⁻/⁻^ female: *n* = 14; at twelve months of age, WT male: *n* = 22, *Kcnj15*^⁻/⁻^ male: *n* = 10, WT female: *n* = 22, *Kcnj15*^⁻/⁻^ female: *n* = 10.

Young *Kcnj15*^⁻/⁻^ males appear to transition more rapidly from a high-alertness state to a self-directed coping response. This interpretation is supported by the observation that when these same mice were reassessed at 12 months of age, latency to first center entry no longer differed significantly among the groups (**Fig. 5e**). These results suggest normalization of initial avoidance behavior with age. Notably, at 12 months of age, *Kcnj15*^⁻/⁻^ males spent longer durations in the center compared to WT males (*P* = 0.0419; **Fig. 5f**), while grooming and rearing behaviors remained comparable across genotypes and sexes (**Fig. 5g–h**). This transition from avoidance at 6 months of age in male mice (**Fig. 5a**) to potential disinhibition or altered risk assessment at 12 months (**Fig. 5e**), suggests dynamic remodeling of limbic circuitry across the lifespan, rather than a static anxiety trait. This pattern is reminiscent of compensatory circuit adaptations or progressive rearrangements in cortico-striatal-limbic networks reported in other genetic models [10, 33]. Loss of Kir4.2 may therefore disrupt the plasticity of circuits regulating emotional reactivity and stress coping, leading to shifting patterns of anxiety-related behavior over the life span rather than a static trait phenotype.

### Loss of *Kcnj15* drives region-specific dopaminergic neurodegeneration, pathological α-synuclein accumulation, and microgliosis

Many widely used α-synA53T and recessive PD models show limited or inconsistent SNpc dopaminergic cell loss over standard timeframes, limiting their utility for testing neurorestorative therapies [5, 7]. To determine whether *Kcnj15* deficiency accurately models the neuropathological hallmarks of PD, we evaluated dopaminergic integrity, abnormal α-synuclein aggregation, and neuroinflammation in the substantia nigra brain area, the epicenter of PD pathology [14], in 18-month-old *Kcnj15*^⁻/⁻^ and WT littermates (**Fig. 6a-6g**). Low-magnification whole-brain coronal sections immunostained for tyrosine hydroxylase (TH, red) and phosphorylated α-synuclein (α-syn-pSer129, green) revealed a macroscopic reduction in TH immunoreactivity within the midbrain dopaminergic nuclei of *Kcnj15*^⁻/⁻^ mice (Fig. 6c) compared to age-matched WT controls (**Fig. 6a**). Enlarged images of these regions confirmed profound, multi-modal pathological changes (**Fig. 6b&6d**). In WT mice, the midbrain exhibited dense TH^+^ neuronal networks, resting IBA1^+^ microglia (red), and minimal pathological α-syn-p signal (**Fig. 6b**). In contrast, *Kcnj15*^⁻/⁻^ mice displayed a visible loss of TH^+^ dopaminergic soma and fibers, accompanied by prominent punctate accumulation of pathological α-syn-p (**Fig. 6d**, right panel). Furthermore, *Kcnj15*^⁻/⁻^mice exhibited marked signs of neuroinflammation. IBA1^+^ microglia displayed reactive morphologies and increased co-expression of the major histocompatibility complex class II marker HLA-DR (green), indicative of active microgliosis (**Fig. 6d**, left panel). Dual-labeling also revealed frequent co-localization of α-syn-p within or closely adjacent to IBA1^+^ microglia (**Fig. 6d**, middle panel), suggesting active microglial engulfment of pathological α-synuclein species.

**Figure 6.**
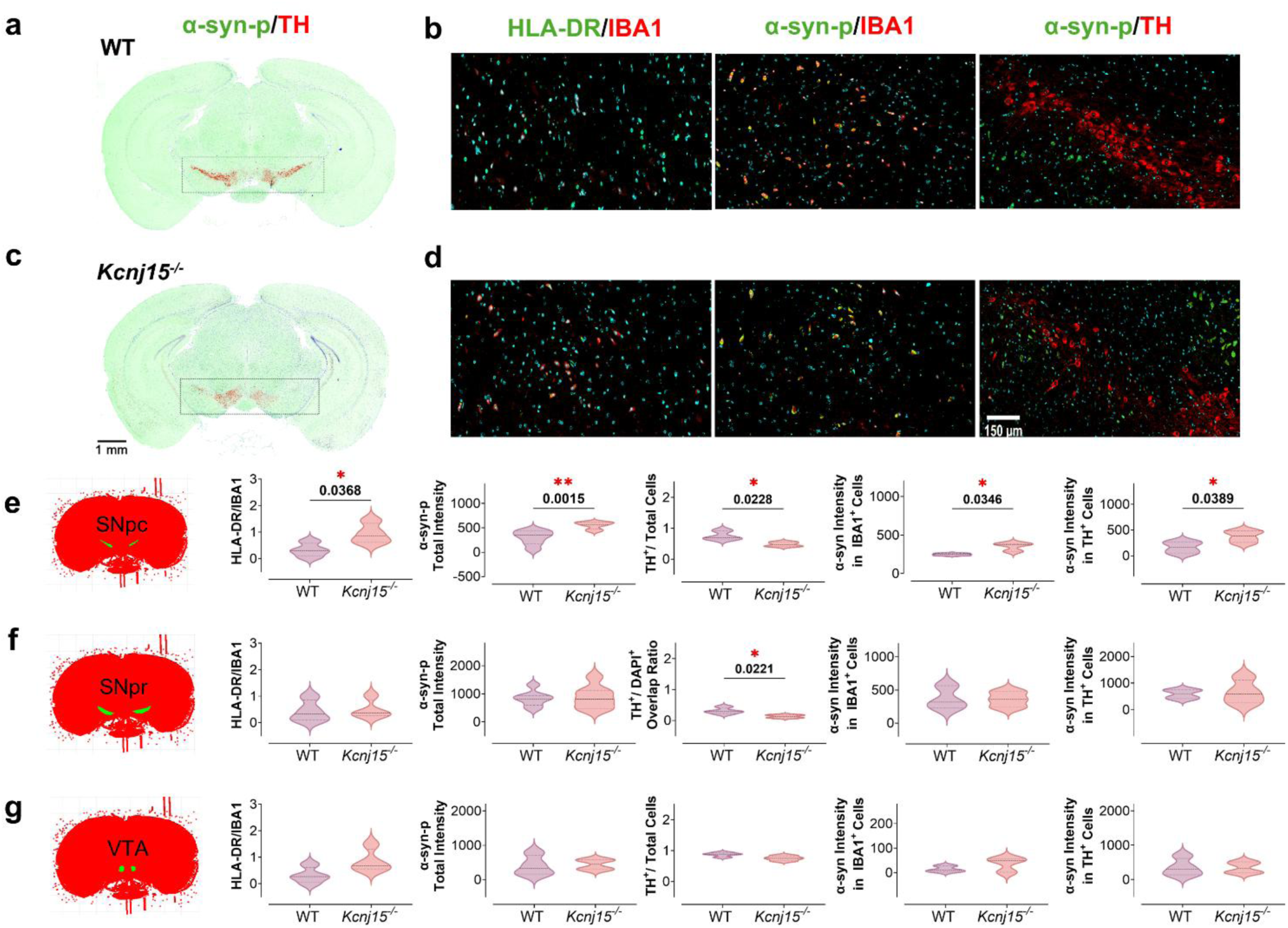
Region-specific microglial activation, phosphorylated α-synuclein pathology, and dopaminergic neuron loss in *Kcnj15*^⁻/⁻^ mouse midbrain. Representative immunohistochemistry (IHC) images (**a-d**) and quantitative analyses of microglial activation, phosphorylated α-synuclein (α-syn-pSer129) pathology, and dopaminergic neuron degeneration in the midbrain of 18-month-old female wild-type (WT) and *Kcnj15* knockout (*Kcnj15^-/-^*) mice (**e-g**). Low-magnification images of coronal brain sections (**a & c**) indicate the anatomical regions of interest used for analysis. High-magnification representative images (**b&d**) show co-immunolabelling for HLA-DR (green) and IBA1 (red) to identify activated microglia, as well as α-syn-pSer129 (green) co-labeled with either IBA1 (red) or tyrosine hydroxylase (TH, red). (**e-g)** Coronal brain sections, aligned with all the slides used for the analysis, encompass the substantia nigra pars compacta (SNpc, **e**), substantia nigra pars reticulata (SNpr, **f**), and ventral tegmental area (VTA, **g**). Violin plots depict quantitative comparisons between WT and *Kcnj15*^⁻/⁻^ mice for each region across five metrics: HLA-DR/IBA1 signal intensity, total α-syn-pSer129 burden, relative dopaminergic survival/density (proportion of TH⁺ neurons in the SNpc and VTA, and the ratio of TH-associated nuclei in the SNpr serving as a proxy for descending dendritic innervation density), α-syn-pSer129 intensity within IBA1⁺ microglia, and α-syn-pSer129 intensity within TH⁺ structures (neuronal somas in SNpc/VTA, and dendritic processes in SNpr), as indicated on individual y-axes. Central lines denote the median with distribution shown by the violin width. Statistical significance between genotypes was assessed using unpaired two-tailed *t*-tests (*n* = 3-7). These data demonstrate region-dependent alterations in microglial activation, α-syn-pSer129 accumulation, and dopaminergic integrity associated with loss of *Kcnj15*.

To rigorously assess these observations, we performed targeted, automated quantification across three distinct midbrain sub-regions (**Fig. 6e-6g**), revealing a pattern of vulnerability that closely mirrors human PD[2]. *Kcnj15*^⁻/⁻^ mice exhibited a significant increase in activated microglia (HLA-DR⁺ IBA1⁺) within the SNpc, the most vulnerable area in PD (*P* = 0.0368, *n* = 4, **Fig. 6e**), sparing the adjacent VTA and substantia nigra pars reticulata (SNpr) (**Fig. 6f & 6g**). Given the established interplay between microglial activation and α-synuclein seeding and aggregation in PD [34], we next examined whether Kir4.2 deficiency alters the accumulation of phosphorylated α-synuclein (α-syn-pSer129), the pathological hallmark of Lewy bodies. Consistent with the microglial hyperactivation, significantly elevated levels of α-syn-pSer129 were observed in the SNpc area (*P* = 0.0015 *n* = 4; **Fig. 6e**, α-syn-p total intensity), but not in the SNpr (**Fig. 6f**) nor VTA areas (**Fig. 6g**). Notably, the increased α-syn-pSer129 accumulation was frequently localized within IBA1^+^ microglia (*P* = 0.0346, *n* = 3, **Fig. 6d & 6e**, α-syn intensity in IBA1^+^ cells). These findings suggest the absence of Kir4.2 enhances microglial uptake of α-syn-pSer129 and/or that there may be impaired clearance of this pathological α-synuclein species. It can be suggested that Kir4.2 deficiency may not only potentiate α-synuclein-induced microglial activation but also promote a state in which microglia act as reservoirs and amplifiers of α-syn pathology. This response may be speculated to establish a feed-forward inflammatory loop.

Crucially, the inflammatory milieu in the absence of Kir4.2 was associated with frank neurodegeneration and structural dopaminergic loss. The *Kcnj15*^⁻/⁻^ mice exhibited a significant reduction in TH⁺ neurons within the SNpc compared with WT animals (*P* = 0.0228, *n* = 3; **Fig. 6e**, TH^+^/total cells). Concurrently, in the adjacent SNpr, which is predominantly innervated by descending SNpc dendrites rather than resident dopaminergic somas [35], there was a significant decrease in dopaminergic fiber density, quantified as a reduced ratio of TH-associated nuclei (*P* = 0.0221, *n* = 4; **Fig. 6f**, TH^+^/DAPI^+^ overlap ratio). Surviving TH⁺ neurons in the SNpc of *Kcnj15*^⁻/⁻^ mice displayed significantly elevated intraneuronal α-syn-pSer129 accumulation (*P* = 0.0272, *n* = 4, **Fig. 6e**, α-syn intensity in TH^+^ cells). This observation is consistent with enhanced pathological α-syn aggregation and/or impaired proteostatic clearance mechanisms resulting from Kir4.2 deficiency. No significant α-syn-pSer129 accumulation nor dopaminergic neuron loss was evident in the VTA, the most degeneration-resilient area among the three (**Fig. 6g**). These results indicate the effects of Kir4.2 loss are anatomically selective.

Collectively, these results demonstrate that Kir4.2 deficiency precipitates a constellation of pathological changes characteristic of early Parkinsonian neurodegeneration. The observed pattern of significant microglial activation, α-syn-pSer129 accumulation, and TH⁺ neuron loss in the SNpc contrasted with a reduction in dopaminergic dendritic innervation in the SNpr and minimal dopaminergic cell loss in the VTA. These observations closely mirror the selective vulnerability of the nigrostriatal pathway observed in human PD [36] and are supported by the behavioral phenotypes observed in the *Kcnj15*^⁻/⁻^ mice (**Figs 1-5**). These data implicate Kir4.2 as a new modulating factor involved in α-syn-associated neurodegeneration.

### Striatal transcriptomics reveals a coherent oligodendrocyte–myelination enrichment signature in *Kcnj15*^⁻/⁻^ mice

Finally, to interrogate the downstream molecular consequences of the nigral dysfunction described above in *Kcnj15*^⁻/⁻^ mice (**Figs 1-6**), we performed bulk RNAseq on the striatum, the primary projection target of SNpc neurons [37], using 18-month-old female mice. While global gene expression changes were generally modest (**Fig. 7a**), as is typical for bulk tissue profiling due to cellular heterogeneity, the genes showing genotype-associated shifts converged on a highly consistent biological theme across independent enrichment approaches.

**Figure 7.**
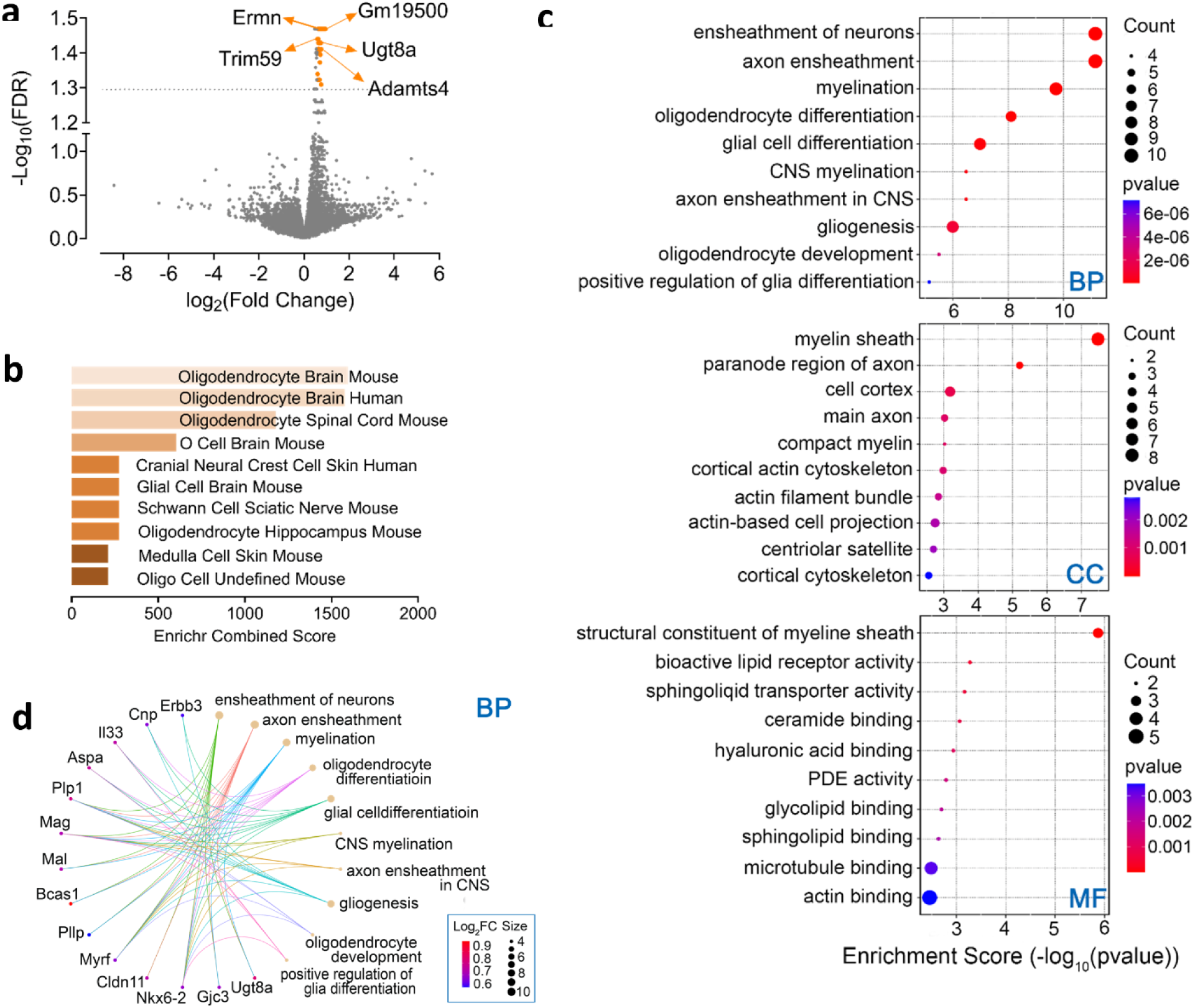
Transcriptomic profiling identifies a robust oligodendrocyte- and myelination-associated signature in the *Kcnj15*^⁻/⁻^ striatum. (a) Volcano plot showing differential gene expression between WT and *Kcnj15*^⁻/⁻^ striatal samples. Each dot represents one gene, plotted as log₂ (Fold Change) versus −log₁₀ (false discovery rate, FDR). Genes passing the selected significance thresholds (|-log_10_FDR| >1.3, |log₂FC| > 0.585) are highlighted in orange, while non-significant genes are shown in grey. The top five most significant DEGs are labelled. Volcano plot was generated using the SRPLOT platform [56]. (**b)** Cell-type enrichment analysis of differentially expressed genes showing the top 10 enriched terms from the CellMarker 2024 gene set library ranked by combined score (*z* x (-Log*P*), where z is the Z-score computed to assess the deviation from the expected rank of that term, *P* is the unadjusted *P*-value calculated using a Fisher’s exact test) in Enrichr [57]. The most significant categories are represented by the longest and brightest bars, revealing that the differentially expressed genes are predominantly enriched for oligodendrocyte-specific signatures across both mouse and human datasets. *Note*: “O Cell” and “Oligo Cell” are nomenclature variants of oligodendrocytes curated by the CellMarker 2024 database from independent transcriptomic studies. **(c)** Gene Ontology (GO) enrichment analysis of differentially expressed genes across the three GO domains: Biological Process (BP), Cellular Component (CC), and Molecular Function (MF). Dot size represents the number of genes associated with each term, while colour indicates enrichment significance (*P*-value). Enriched terms highlight processes related to axon ensheathment, myelination, oligodendrocyte differentiation, cytoskeletal organisation, and lipid-binding functions. **(d)** Gene-concept network (cnet) plot illustrating the relationships between differentially expressed genes and enriched GO terms for the BP category. GO terms are shown on the right, genes on the left, and edges indicate gene-term associations. Node size reflects gene count per term, and gene node colour represents log₂ (fold change), revealing shared driver genes underlying oligodendrocyte maturation, myelin structure, and axonal support pathways.

Cell-type enrichment analysis against reference signatures showed the strongest enrichment for oligodendrocyte-lineage annotations, including “Oligodendrocyte Brain Mouse” and “Oligodendrocyte Brain Human,” with additional enrichment across related glial a nd myelinating cell annotations (**Fig. 7b**). The striatal transcriptional changes associated with *Kcnj15* loss are therefore disproportionately aligned with oligodendrocyte biology rather than broadly distributed across the neuronal system.

Gene Ontology (GO) enrichment analysis further supported this interpretation (**Fig. 7c**). In the Biological Process (BP) category, the top-enriched terms were dominated by axon ensheathment and myelination, including “ensheathment of neurons,” “axon ensheathment,” “myelination,” and multiple oligodendrocyte differentiation/development terms ( **Fig. 7c**, BP). In the Cellular Component (CC), enrichment was concentrated in myelin- and axon-associated compartments, including “myelin sheath,” “compact myelin,” and “paranode region of axon,” alongside cytoskeletal localisations (**Fig. 7c**, CC). In the Molecular Function (MF), enriched terms included “structural constituent of myelin sheath” and lipid-binding/transport-related functions (*e.g.*, sphingolipid and glycolipid binding/transport), consistent with myelin membrane composition and maintenance programs (**Fig. 7c**, MF). These GO enrichment findings align with the cell type enrichment analysis, reflecting the specialized role of oligodendrocytes as the primary myelinating glia of the central nervous system. These cells are fundamentally responsible for the assembly of the myelin sheath, a multilamellar, lipid-rich membrane that encircles axons [38].

To visualize the gene drivers underlying these enriched categories, we analyzed the gene-concept network (cnet). These networks demonstrated that multiple enriched GO terms are supported by overlapping gene sets, indicating a shared core mechanism. Notably, BP (**Fig. 7d**) and CC (**Supplementary Figure 8a**) networks highlighted canonical oligodendrocyte and myelin-associated genes (*e.g.*, *Plp1, Cnp, Mag, Mal, Bcas1, Pllp, Cldn11, Myrf, Nkx6-2*). In contrast, MF enrichment captured functions consistent with myelin structural components and lipid-related molecular activities (**Supplementary Figure 8b**). Collectively, these results suggest that *Kcnj15* loss in the striatum may be associated with the oligodendrocyte differentiation/maturation and the axon-myelin structural axis.

## Discussion

Despite decades of research, a major bottleneck in PD drug development remains the lack of highly translatable mouse models exhibiting consistent, spontaneous nigrostriatal neurodegeneration for testing neurorestorative therapies. Existing experimental models often rely on massive exogenous stressors or extreme viral overexpression, frequently failing to capture the spontaneous, progressive, and anatomically selective nature of human PD. Widely utilized transgenic models overexpressing human mutant α-synuclein (e.g., A53T, A30P) successfully recapitulate widespread protein aggregation; however, they predominantly develop severe spinal cord and brainstem pathology leading to early paralysis, rather than the anatomically progressive, SNpc-specific dopaminergic neurodegeneration characteristic of human PD [3–5]. Similarly, standard knockout models of autosomal recessive PD genes (e.g., *PINK1*, *PRKN*, *DJ-1*) fail to exhibit spontaneous nigrostriatal dopaminergic cell loss or Lewy-like α-synuclein pathology over standard experimental timeframes, requiring the artificial introduction of massive exogenous stressors to provoke a phenotype [6, 7]. Consequently, establishing an endogenous genetic model that naturally mirrors human pathophysiological logic remains a critical, unmet preclinical need.

By generating and characterizing the *Kcnj15*^⁻/⁻^ mouse, which models a loss-of-function dominant-negative variant identified in familial PD, we establish the inwardly rectifying potassium channel Kir4.2 as a functional guardian of nigrostriatal circuit integrity [8, 9]. Our findings demonstrate that Kir4.2 deficiency is sufficient to drive a triad of PD-relevant pathologies: **(1)** a progressive “coordination-first” motor syndrome with specific cognitive and anxiety-related phenotypes; **(2)** an anatomically selective nigral neuroimmune axis characterized by microglial activation, synucleinopathy, and dopaminergic cell loss; and **(3)** a striatal transcriptional response dominated by oligodendrocyte maturation and myelin remodeling. These findings close the loop on prior genetic identification of *KCNJ15* as a familial PD locus and provide mechanistic support for the Kir4.2 channel as a new biological target contributing to PD susceptibility.

The behavioral trajectory of the *Kcnj15*^⁻/⁻^ mouse offers a nuanced recapitulation of prodromal and early-stage clinical PD. Rather than a monolithic motor failure, these mice exhibit early deficits in dynamic balance and fine motor coordination that precede broader declines in spontaneous locomotion by months, while spatiotemporal gait metrics assessed by DigiGait remain largely preserved. This dissociation mirrors the human clinical picture, where postural instability and complex motor integration deficits frequently manifest before the characteristic shuffling gait, which requires more extensive nigrostriatal compromise [14, 20, 39]. The preservation of gait parameters, typically sensitive to spinal and brainstem circuit dysfunction or advanced asymmetric nigrostriatal degeneration [22, 40], supports the interpretation that Kir4.2 loss first disrupts higher-order basal ganglia–cerebellar motor integration, leaving central pattern generators intact until later stages.

Alongside these motor features, Kir4.2 deficiency drives age-dependent non-motor symptoms, including progressive long-term spatial memory deficits and dynamic shifts in anxiety-like behaviors. This selective impairment of remote memory consolidation is consistent with the particular vulnerability of extended hippocampal–cortical and frontostriatal networks to neurodegeneration and mirrors the insidious executive and memory decline recognized as a core non-motor feature of PD [41, 42]. Furthermore, the *Kcnj15*^⁻/⁻^ mice exhibited a transition from early center-avoidance and accelerated grooming onset to later disinhibition or altered risk assessment in the open field test. This phenotype is consistent with compensatory remodelling or progressive re-weighting within cortico–striatal–limbic circuits governing threat appraisal and coping strategy selection, and parallels the dynamic anxiety phenotypes increasingly recognized in prodromal PD [14, 43, 44].

The most mechanistically convergent finding of this study is the profound anatomical selectivity of neurodegenerative and neuroinflammatory response, which closely recapitulates the unique topography of human PD. Unlike many existing genetic models that exhibit widespread, non-specific pathology, Kir4.2 deficiency precipitates frank dopaminergic neuron loss strictly within the SNpc, with accompanying reduction of descending dopaminergic dendritic innervation in the SNpr, while the VTA is entirely spared. This regional vulnerability is tightly coupled to a localized neuroimmune response characterized by profound microglial hyperactivation and the accumulation of phosphorylated α-synuclein (α-syn-pSer129). The frequent co-localization of α-syn-pSer129 within reactive microglia suggests a state of “frustrated phagocytosis,” wherein microglia engulf pathological protein species but fail to efficiently degrade them, becoming cellular reservoirs that seed further aggregation and amplify neuroinflammation [45].

Why does Kir4.2 loss produce this specific pathology? We propose that Kir4.2 deficiency shifts microglia from effective homeostatic surveillance toward a dysfunctional, chronically activated state. In the absence of functional Kir4.2, microglia may undergo a maladaptive metabolic shift, compromising their clearance capacity and transforming them into cellular reservoirs that amplify α-synuclein seeding. The strict confinement of this pathology to the SNpc aligns with evidence that SNpc microglia maintain intrinsically distinct transcriptomic profiles and baseline activation thresholds relative to other brain regions [46, 47], suggesting that Kir4.2 may be particularly critical for constraining neuroimmune set-points within this vulnerable midbrain niche. Intraneuronal α-syn-pSer129 accumulation in surviving TH⁺ neurons further implies concurrent proteostatic failure at the neuronal level, consistent with impaired proteasomal or autophagic clearance mechanisms downstream of Kir4.2 loss.

Our striatal transcriptomic analysis introduces an additional, underappreciated axis to the Kir4.2 phenotype. Although global differential expression was modest, which is typical for bulk tissue profiling given cellular heterogeneity, the genes showing genotype-associated shifts converged robustly on oligodendrocyte maturation and myelination biology, supported by enrichment of canonical myelin-lineage genes including Plp1, Mag, Cnp, Mal, Bcas1, Cldn11, Myrf, and Nkx6-2.

While frequently overlooked in classic models of PD, oligodendrocytes are metabolically coupled to axons and dynamically regulate conduction velocity, an essential parameter for the temporally precise integration of basal ganglia computations [48]. Emerging multi-omics and single-cell studies report oligodendrocyte lineage perturbations and myelin support pathway alterations in human PD tissue [49]. Upregulation of myelination genes in the *Kcnj15⁻/⁻* striatum may initially represent an adaptive, compensatory response to early dopaminergic axonal stress, attempting to bolster metabolic support and ensheathment [50]. However, spatially biased or excessive myelin remodelling can become maladaptive. By retuning conduction velocities and spike-timing relationships across long-range axons, it risks shifting network synchrony toward the rigid, hyper-oscillatory operating regimes characteristic of the Parkinsonian state [18, 51]. Furthermore, a persistent shift toward mature myelinating states depletes the oligodendrocyte precursor pool, constraining the experience-dependent myelin plasticity required for adaptive motor learning and cognitive flexibility [52]. Bidirectional cross-talk between the oligodendrocyte lineage and activated microglia likely further amplifies this dysfunction, with inflammatory cues and myelin-related signals exacerbating one another in a feed-forward cycle [53]. Protein-level validation and future single-nucleus or spatial transcriptomic studies will be required to determine whether this striatal signature reflects altered oligodendrocyte cell numbers, per-cell transcriptional dysregulation, or both.

An attractive working model therefore emerges in which Kir4.2 deficiency drives PD-relevant pathology through two coordinated, mutually reinforcing axes (**Fig. 8**): a nigral neuroinflammatory–synucleinopathy axis in which microglial dysfunction, α-synuclein seeding, and dopaminergic neuron loss progressively degrade nigrostriatal signalling; and a striatal oligodendrocyte–myelin axis in which altered conduction timing and reduced axonal metabolic support lower the threshold for network dysfunction and neurodegeneration. Rather than acting in isolation, these pathways likely exacerbate one another. Dopaminergic dysfunction and neuroinflammation reshape glial states, while myelin dysregulation amplifies circuit instability and accelerates degeneration, collectively driving the age-dependent progression of the PD-like phenotype observed here. These findings establish the *Kcnj15⁻/⁻* mouse as a uniquely robust, endogenous genetic model for prodromal and early-stage PD and identify Kir4.2-dependent neuroimmune and glial signalling as a compelling therapeutic axis for disrupting the cycle of inflammation and neurodegeneration before irreversible circuit collapse occurs.

**Figure 8.**
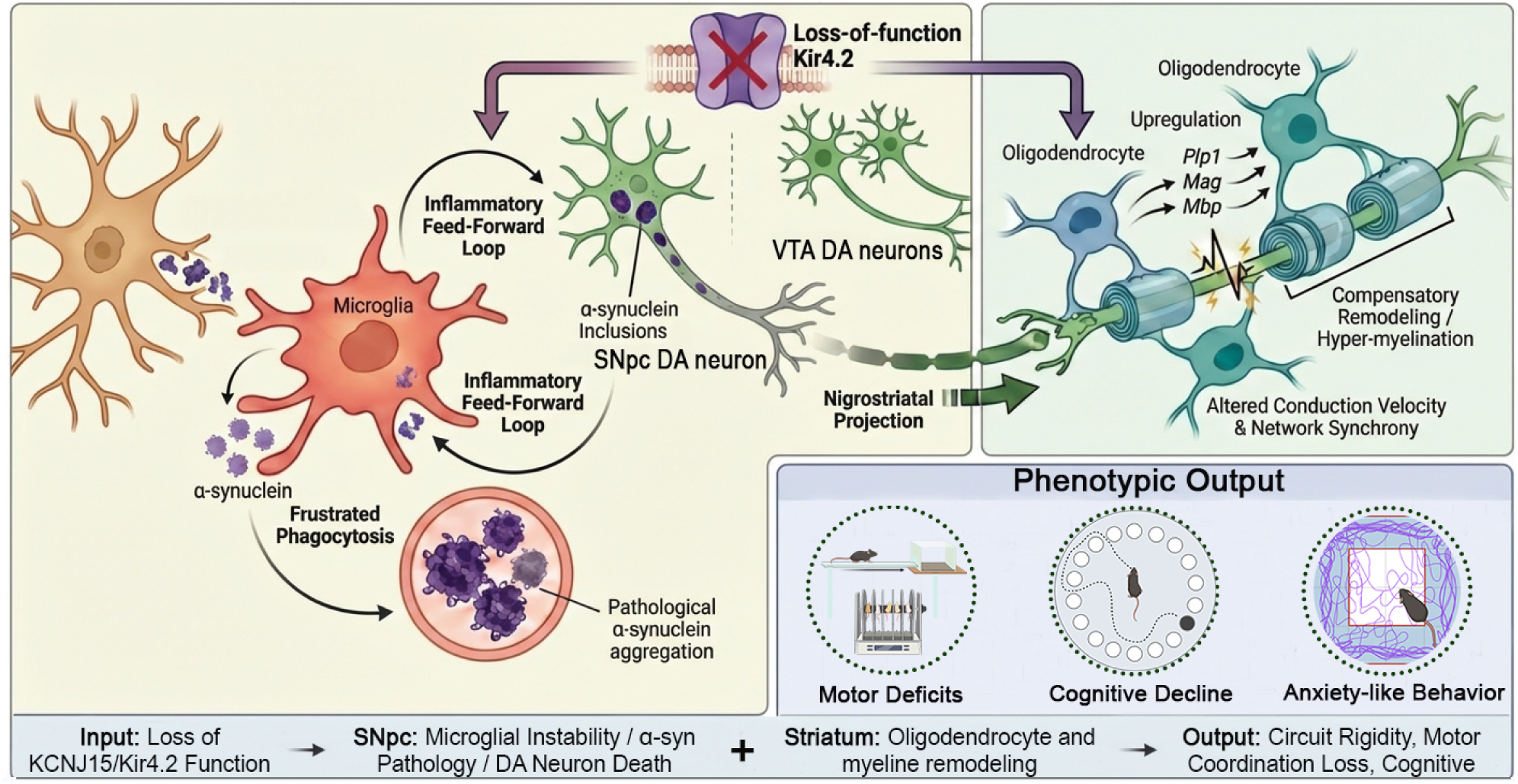
Schematic overview of a proposed working model linking loss of Kir4.2 function to Parkinson’s disease–related pathology via coordinated glial and neuronal dysfunction. In this hypothesized model, global Kir4.2 deficiency creates a vulnerable nigrostriatal environment characterized by coordinated glial and neuronal dysfunction. Based on our histological observations, we propose that Kir4.2 loss may directly or indirectly promote microglial instability. This could potentially impair intracellular α-synuclein clearance (*e.g.*, via frustrated phagocytosis), contributing to a self-amplifying inflammatory feed-forward loop. This neuroinflammatory milieu and accumulation of α-synuclein inclusions are associated with profound dopaminergic (DA) neuronal stress and degeneration in the substantia nigra pars compacta (SNpc), while sparing DA neurons in the ventral tegmental area (VTA). Concurrently, transcriptomic profiling suggests that Kir4.2 dysfunction affects the nigrostriatal circuitry by engaging oligodendrocyte stress responses in the striatum. The latter is marked by a compensatory up-regulation of myelin-associated genes (*e.g.*, *Plp1*, *Mag*, *Mbp*), and potential maladaptive axonal remodeling. We propose that these interacting, multi-cellular changes disrupt conduction velocity and network synchrony, collectively giving rise to PD-relevant phenotypes such as motor deficits, cognitive decline, and anxiety-like behaviors.

## Methods

### Animals and Housing Conditions

Heterozygous *Kcnj15*^⁻/⁻^ C57BL6/J mice were generated by the Monash Genome Modification Platform (Melbourne) and handled in accordance with *the Australian Code for the Care and Use of Animals for Scientific Purposes*. All procedures were approved by the Griffith University Animal Ethics Committee (GRIDD/02/22/AEC). Homozygous WT and *Kcnj15*^⁻/⁻^ mice were selectively bred from these heterozygous mice, with genotype confirmation by PCR using the Phire Tissue Direct PCR kit (Thermo Fisher Scientific). The mice were housed in individually ventilated cages with well-designed structural features that allowed hiding and environmental enrichment, including tissues, tunnels, nesting material, and chew sticks. Animals had *ad libitum* access to dry food and water in a well-ventilated room maintained at 22 ± 2 °C with 55 ± 10% relative humidity, under a 12-hour light-dark cycle (lights on from 7:00 AM to 7:00 PM) facilitated by the animal facility staff at Griffith University.

### Animal Behavioral Tests

Before beginning any behavioral tests, the mice were acclimated to the behavioral study room for 1 h, with background music playing at a volume of ∼62-65 dB and lighting set between 250-400 lux. To minimize anxiety further, non-aversive handling techniques, such as acclimation to the handlers and tunnel handling, were employed during the tests.

### The Open Field Test of Locomotion and Anxiety

The open-field arena was a white square enclosure (60 x 60 cm; wall height 40 cm). Each mouse was recorded using a GoPro Hero11 (GoPro) while freely exploring the arena for 10 min. Mouse behavior was analyzed using EthoVision XT17 video tracking software (Noldus), which overlays user-defined zones onto the video and tracks locomotion and location over time. For analysis, the arena was divided into two configurations: one with 16 equal areas (Rows A-D, Columns 1-4), and another with four zones (outer, edge, middle, and center) (**Supplementary Fig. 1**). This design allows for detailed analysis of various parameters, including velocity and travel distance within specific areas or zones, line crossings (horizontal and vertical beam breaks), time spent mobile or stationary in each area/zone, and the location and frequency of grooming and rearing behaviors.

### The Barnes Maze Test of Spatial Memory and Learning

The Barnes maze test consists of a circular platform (120 cm diameter) with 40 evenly spaced holes around the perimeter, one of which led to a black hiding/escape box. Overhead illumination served as an aversive stimulus to motivate exploration and escape. Visual cues were positioned around the room to support spatial navigation. At the start of each trial, each mouse was placed in the center of the arena under an opaque cover for 1 min with the light off. After the light was turned on and the cover removed, the mouse was allowed to explore the maze for 3 min, while being recorded with a GoPro camera.

Mice that failed to locate the escape box within the allotted training time were gently guided to the target hole and allowed to remain in the box for 1 min before being returned to their home cages. Between trials, the platform was cleaned with 80% ethanol and rotated to maintain the escape hole’s spatial position in the room while minimizing odor cues. Training was performed for four consecutive days. A probe trial on day 5 was conducted with the escape box removed to assess short-term memory, and a second probe trial on day 12 assessed long-term memory under the same conditions. The recordings were analyzed using EthoVision XT17.

### Accelerating Rotarod Test of General Motor Function

General motor performance, *i.e*., balance and coordination as well as strength and endurance, was assessed using the Rotarod system (Ugo Basile). Mice were first placed on the rod rotating at 4 rpm to acclimate, after which the program accelerated from 4 rpm to 60 rpm over 300 s. Following the acceleration phase, the rod continued at 60 rpm until the mice fell, triggering automatic recording of the event. Each mouse completed three trials separated by a 15 -min inter-trial interval. The latency to fall and total distance travelled were exported for downstream analysis.

### The Balance Beam Test of Balance and Coordination

The balance beam test assesses balance and coordination in rodents, requiring mice to traverse a 5 mm-thick acrylic beam (75 cm long) to reach an escape box at the opposite end. Mice were first acclimated to the escape box for 2 min before being placed on the beam at the end furthest from the box. The mice then crossed the beam to reach the box, after which they were allowed to rest for 1 min before repeating the test. Each mouse completed three technical replicates *per* day across five consecutive days (four training days followed by one test day). Recordings were analyzed to measure the traversal time and the number of foot slips, defined as instances when a toe slipped off the top surface of the beam before the mouse regained balance.

### DigiGait Gait Analysis

Gait was assessed using the DigiGait system (Mouse Specifics), which consists of an enclosed treadmill with a transparent belt and a high-speed camera positioned beneath the belt recording at 147 frames/s to capture a ventral view of the mouse as it runs. DigiGait Analysis software was used to quantify spatiotemporal gait parameters. For this test, the mice underwent a 5-day training period to ensure consistent results. On day 1, mice were placed in the enclosed chamber and allowed to acclimate until rearing and grooming behaviors were observed, with the acclimation time recorded. Each mouse was recorded running at 15 cm/s for 30 s, with 1-min rest intervals between replicates (3 replicates). On days 2 to 5, after the acclimation period, each mouse was recorded running at 15 cm/s, 20 cm/s, and 25 cm/s for 30 s each. For each run, videos were trimmed to the best 10 s of continuous, steady locomotion (excluding turning or disruptions), and the best segment at each speed was selected for downstream analysis.

### Cardiac Perfusion and Brain Histology Preparation

For immunohistochemistry (IHC), 18-month-old female mice were deeply anaesthetized with 1.5% sodium pentobarbital (60 mg/kg, *i.p.*). After loss of reflexes, a thoracotomy was performed, and a 27G perfusion needle was inserted into the left ventricle of the heart. The right atrium was then incised, and the mouse was perfused with 20 mL of PBS at 200 µL/s to clear blood from the circulatory system, followed by 40 mL of 4% paraformaldehyde (PFA) at the same rate. The brain was carefully removed and post-fixed in 4% PFA for 24 h before being transferred to PBS until sectioning. Paraffin-embedded samples were then sectioned at 5 µm thickness (Translational Research Institute, Brisbane) targeting the substantia nigra pars compacta (SNpc) (AP -3.0 mm to -3.5 mm from bregma).

### Immunohistochemistry Analysis

Paraffin-embedded brain sections were preheated at 56 °C for 15 min, deparaffinised in xylene (2 × 10 min), and rehydrated through graded ethanol washes (100%, 95%, 85%, 75%, and 50% ethanol; 5 min each) following a 1:1 xylene/ethanol step. Slides were then rinsed briefly in cold reverse-osmosis water and incubated in PBS for 30 min. Antigen retrieval was performed by heating slides in 10 mM Tris-Base, 1 mM EDTA buffer (pH 8.0) at 100 °C/20 min, followed by a 10 min rinse under cold running water. Sections were washed twice in TBS-T buffer (0.1% Tween-20 in TBS; 5 min each) and blocked for 2 h at room temperature in 10% goat serum and 1% BSA in TBS-T in a humidified chamber.

Slides were then incubated overnight at 4 °C with the primary antibodies: chicken anti-tyrosine hydroxylase (TH) polyclonal antibody (Abcam, ab76442, 1:1000), chicken anti-IBA1 polyclonal antibody (Thermo Fisher Scientific, PA5-143572, 1:1000), rabbit anti-α-synuclein (phospho-Ser129) polyclonal antibody (Abcam, ab59264, 1:200), and rabbit anti-HLA-DR monoclonal antibody (Thermo Fisher Scientific, MA5-32232, 1:250).

After two TBS-T washes (5 min each), sections were incubated for 1.5 h at room temperature with secondary antibodies: goat anti-chicken IgY Alexa Fluor 647 (Thermo Fisher Scientific, A-21449, 1:500) and goat anti-rabbit IgG Alexa Fluor 488 (Thermo Fisher Scientific, A-11008, 1:500) in TBS-T containing 1% BSA. Sections were washed twice in TBS-T (5 min each), counterstained with DAPI (1 µg/mL; Thermo Fisher Scientific, SRP-62248) for 10 min, and washed again twice before dehydration through graded ethanol and xylene washes in reverse order.

Slides were mounted with coverslips and cured for 24 h before imaging on an Olympus VS200 slide scanner at 20× magnification. Using excitation/emission settings of 360/460 nm (DAPI), 490/525 nm (Alexa 488), and 594/633 nm (Alexa 647).

### RNAseq Profiling of Mouse Striatum Samples

Female WT and *Kcnj15*^⁻/⁻^ (KO) mice (*n* = 5 per genotype; ∼1.5 years old) were euthanized by cervical dislocation. Brains were harvested immediately, snap-frozen in liquid nitrogen, and stored at -80°C. To preserve RNA integrity during processing, brains were immersed in the RNAlater™-ICE solution (Thermo Fisher Scientific) pre-cooled to -80°C and gradually warmed to -20 °C over 24 h. Striatal tissue was then dissected on dry ice and subsequently stored in RNAlater™-ICE solution at -20 °C. RNA extraction from the isolated striatum, library generation, and mRNA sequencing were performed by the Australian Genome Research Facility (AGRF, Brisbane, Australia).

Libraries from 10 samples (5 KO and 5 WT) were sequenced on an Illumina NovaSeq X Plus platform configured for 150 bp paired-end (PE) reads. The sequencing run yielded an average sequencing depth of approximately 36 million paired reads per sample (range: 31.3-40.8 million paired reads). Primary sequence data demultiplexing and FASTQ generation were executed using the Illumina DRAGEN BCL Convert pipeline (v07.031.732.4.3.6). Subsequent quality control and read trimming were performed prior to alignment.

Cleaned paired-end reads were aligned to the mouse reference genome (build *Mus musculus* mm39/GRCm39) using the STAR aligner (v2.3.5a). Mapping was highly robust, with >80% of reads successfully mapping to the reference genome across all samples. Reference-guided transcript assembly was optionally generated using StringTie (v2.1.4) referencing the RefSeq GRCm39 annotation (GCF_000001635.27).

Read quantification at the gene level was performed using featureCounts (Subread v1.5.3). Differential gene expression analysis was subsequently carried out using edgeR (v4.0.9) in R (v4.3.1)[54]. To normalize for varying sequencing depths and composition biases across libraries, the Trimmed Mean of M-values (TMM) normalization method was applied. Finally, generalized linear model (GLM) framework was used to quantify differential expression between the KO and WT groups.

### Data Analysis

Open field, Barnes maze, and balance beam videos were captured using a GoPro Hero11 camera. The Open field and Barnes maze data were analyzed using EthoVision XT17 (Noldus), while the balance beam outcomes were manually scored. DigiGait data were collected and analysed using the DigiGait Analysis software (Mouse Specifics). Rotarod data were exported from the Rotarod system (Ugo Basile) and processed in GraphPad 10.6.0. Fluorescence images quantification was batch-analyzed with Python, CellProfiler and RStudio with scripts available at Github [55]. Datasets were first assessed for normality, then analyzed using Student’s *t*-test, paired *t*-test, one-way ANOVA with Tukey’s multiple comparison test, and simple or multiple linear regression tests, as appropriate. Non-normally distributed data were analyzed using the Friedman test with Dunn’s multiple comparison tests in GraphPad 10.6.0. All data are presented as mean ± SD with biological repeats (*n* number) reported.

## Declarations

### Ethics approval

All experiments were conducted with ethics approval from the Griffith University Animal Ethics Committee, in accordance with the Australian Code for the Care and Use of Animals for Scientific Purposes, protocol no. GRIDD/02/22/AEC.

### Availability of data and materials

All data generated and analyzed during this study are included in this article or available from the corresponding author on reasonable request.

### Consent for publication

All authors have read the final draft of the manuscript and approved its submission.

### Author contributions

B.Garland: Conceptualization and investigation, Writing – original draft, review & editing. Z.S.: Conceptualization and investigation, Writing – review & editing. M.C.: Image analysis of IHC results. B.Gao: Conceptualization and methodology, Writing – review & editing. K.M.: Conceptualization and methodology, Writing – review & editing. D.R.R: Conceptualization and supervision, Writing – review & editing. G.D.M.: Conceptualization, investigation and supervision, Writing – review & editing. L.M.: Conceptualization, investigation, and supervision, Writing – original draft, review & editing. BioRender, Photoshop and Nano Banana were used to generate/edit **Figure 8**; authors take full responsibility for accuracy.

