## Supplementary Figures and Legends for "Kir4.2 deficiency drives progressive Parkinson’s disease-like motor, cognitive and neuropathological phenotypes in mice"

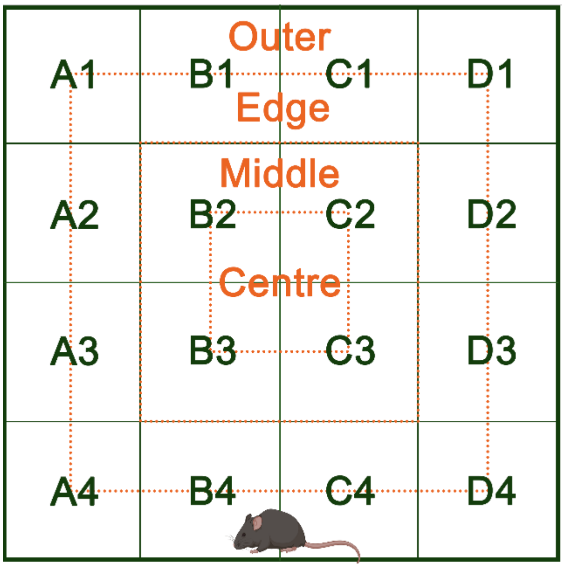


**Supplementary Figure 1. Arena layout of the open field test.** Areas and zones were generated as an overlay using EthoVision XT17 to access locomotion and rearing/grooming behaviour quantitatively. The area was divided into sixteen equal areas (A1-D4) with four separate zones defined (outer, edge, middle, centre).

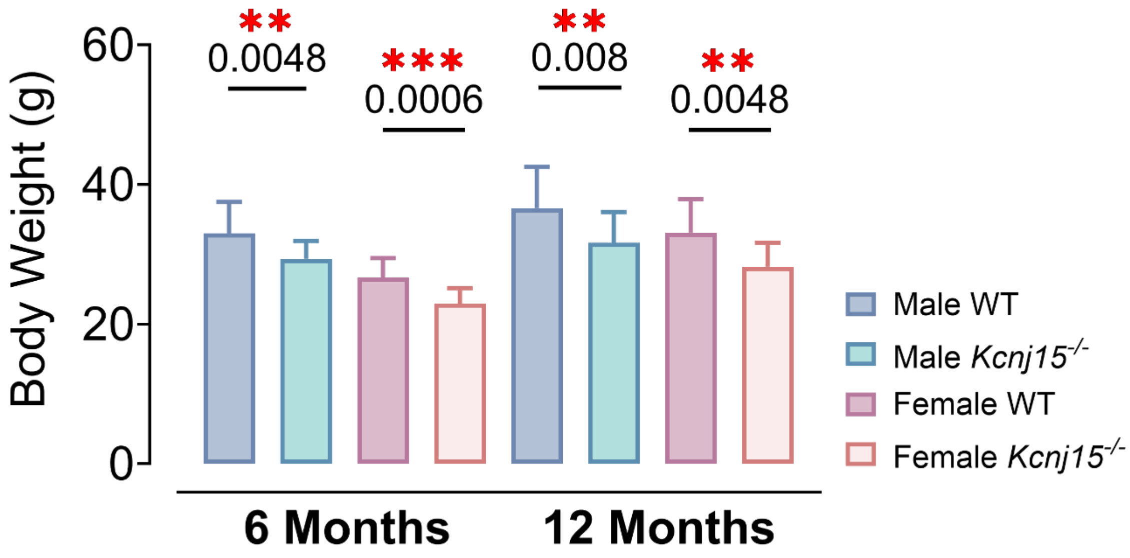


**Supplementary Figure 2. *Kcnj15*$^{-/-}$ mice exhibit significantly reduced body weight compared to their WT controls at both 6 and 12 months of age**. Data are presented as mean ± SD and analysed by unpaired two-tailed *t*-tests. At six months of age, WT male: n = 32, *Kcnj15*$^{-/-}$ male: n = 16, WT female: n = 26, *Kcnj15*$^{-/-}$ female: n = 10; at twelve months of age, WT male: n = 29, *Kcnj15*$^{-/-}$ male: n = 14, WT female: n = 24, *Kcnj15*$^{-/-}$ female: n = 12.

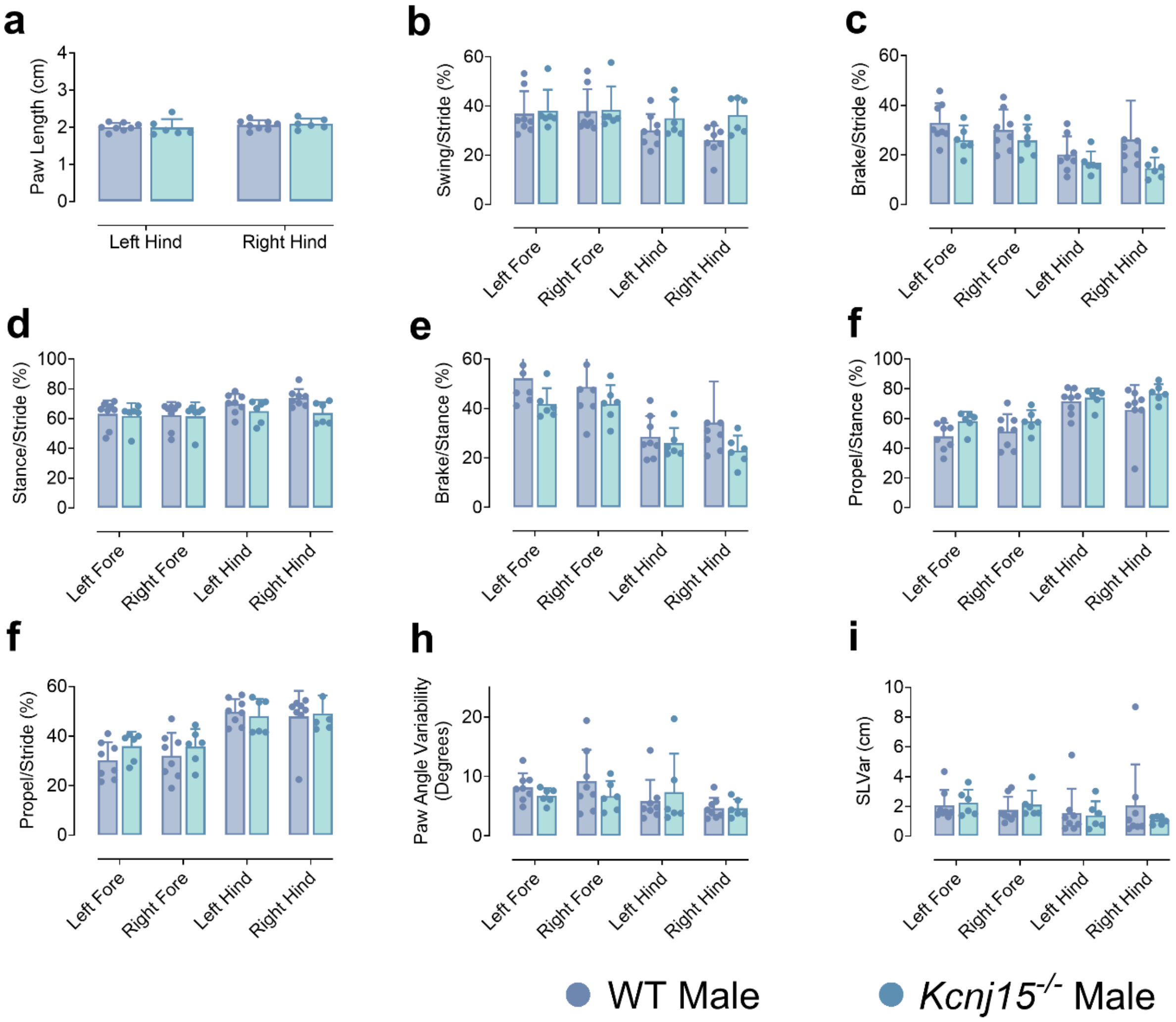


**Supplementary Figure 3. DigiGait reveals preserved limb kinematics and stride-phase proportions in male *Kcnj15*$^{-/-}$ mice.** Spatiotemporal gait parameters were quantified using the DigiGait treadmill system and DigiGait Analysis software in male WT (blue) and *Kcnj15*$^{-/-}$ (magenta) mice. A single stride is composed of two primary phases: the stance phase, during which the paw maintains contact with the treadmill surface, and the swing phase, which transitions into the next step. The stance phase is further subdivided into the brake phase (paw contact increases) and propulsion phase (paw pushes off to propel forward). Paw Angle Variability (degrees) measures how much the paw angle (toe-in/toe-out orientation relative to the direction of travel) varies from step to step. Higher values represent less consistent paw rotation/placement. SLV Var (cm) is stride length variability, which quantifies how much stride length fluctuates across strides. Higher values indicate less consistent stride length from step to step.

Each dot represents an individual mouse. Data were analyzed by unpaired two-tailed *t*-tests and are presented as mean ± SD. Overall, DigiGait metrics were largely comparable between genotypes across limbs, indicating no overt gait-pattern disruption in male *Kcnj15*$^{-/-}$ mice under the tested conditions.

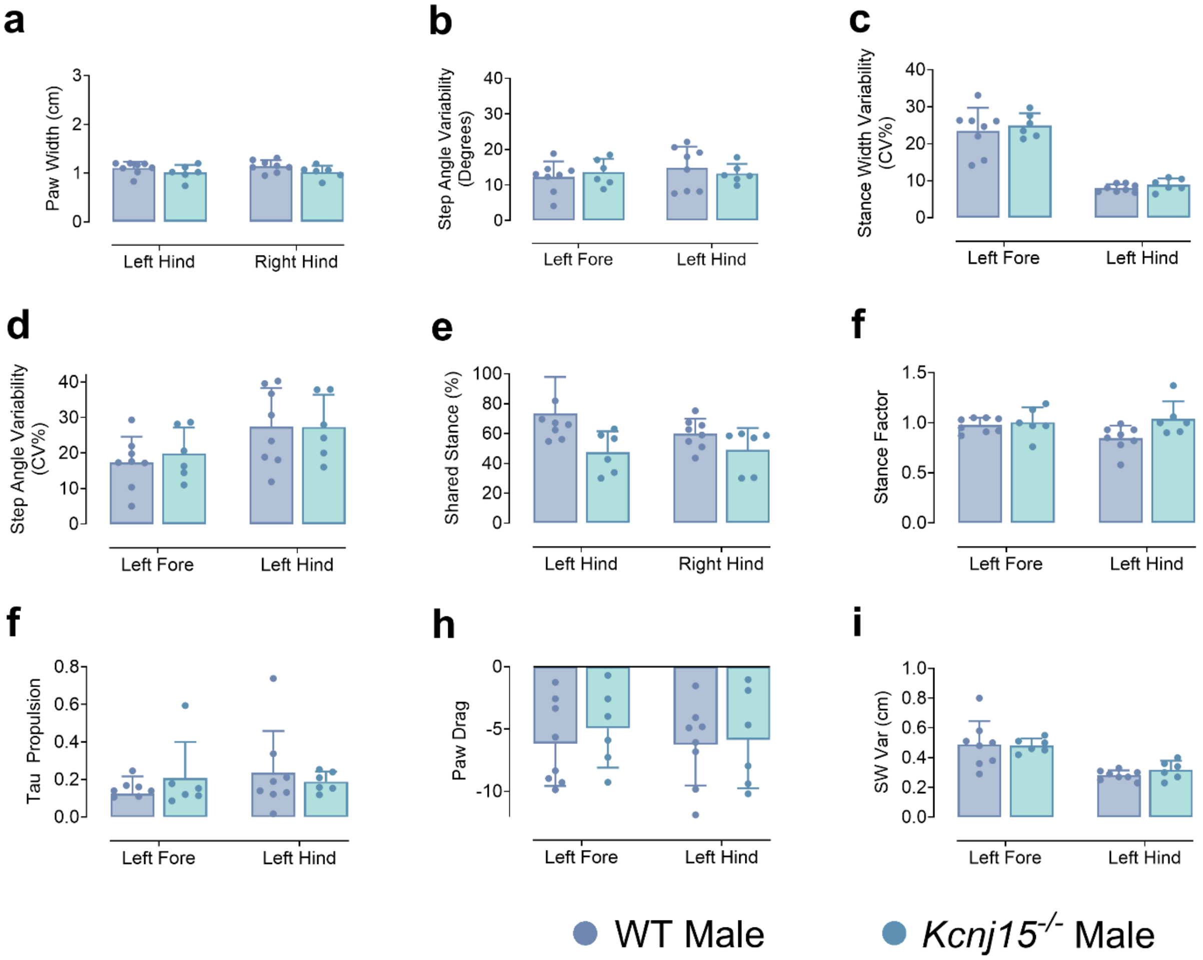


**Supplementary Figure 4. DigiGait analysis of spatiotemporal gait parameters in male WT and *Kcnj15*$^{-/-}$ mice.** Step angle variability measures how much the paw angle (toe-in/toe-out orientation relative to the direction of travel) changes from step to step, expressed as an absolute angular variability in degrees (i.e., more variable paw rotation/placement = higher value). Stance width variability (CV%) quantifies variability stance width (the mediolateral spacing of paw placement / base of support) across steps, expressed as CV%. Shared stance (%) is the percentage of the stride cycle during which the paired limbs are simultaneously in stance. Higher values typically indicate more time with overlapping support. Stance factor is the fraction of the stride spent in stance. Values closer to 1 mean proportionally more time on the belt; smaller values mean proportionally more time in swing. Tau (τ) Propulsion is the “time-constant” descriptor for the propulsion phase of stance (late stance/push-off), reflecting how quickly the paw’s stance/propulsion waveform changes during push-off. Larger τ generally indicates a more gradual propulsion profile; smaller τ suggests a sharper/faster propulsion transition. SW Var (cm) quantifies stride-width variability, which is the variability of mediolateral paw placement / base-of-support across strides. Higher values indicate less consistent stride width from step to step. Each dot represents an individual mouse. Data were analyzed by unpaired two-tailed *t*-tests and are presented as mean ± SD.

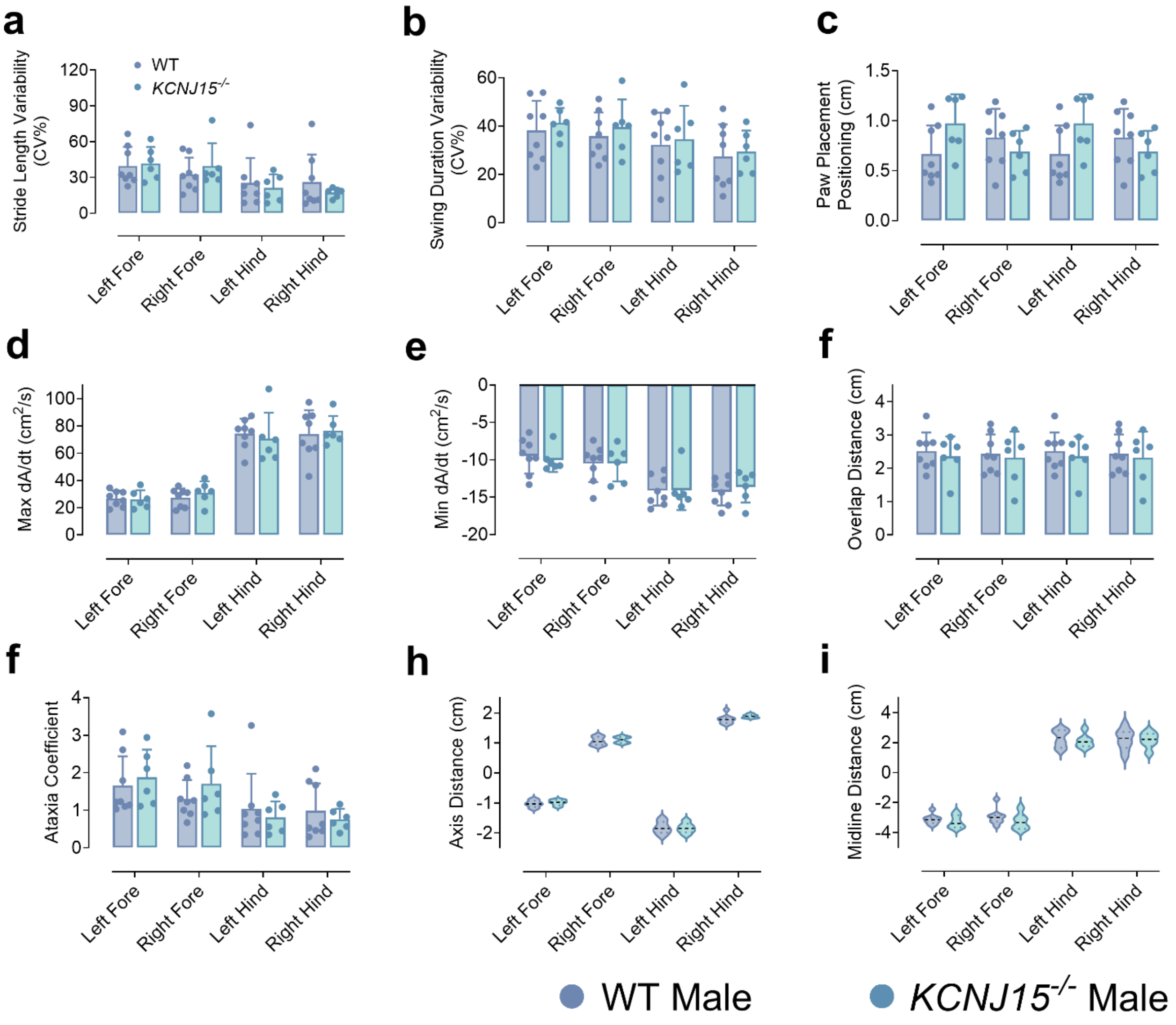


**Supplementary Figure 5. DigiGait analysis of stride variability and paw placement dynamics in male WT and *Kcnj15*$^{-/-}$ mice.** DigiGait treadmill recordings were used to quantify spatiotemporal and paw-contact parameters in wild-type (WT; blue) and *Kcnj15*$^{-/-}$ (magenta) mice. For variability measures, CV% (coefficient of variation) is calculated as *(SD/mean)* × *100* across strides, where higher values indicate greater step-to-step variability. dA/dt represents the time-derivative of paw contact area (A) during stance, reflecting the rate of paw loading (Max dA/dt) and rate of paw unloading (Min dA/dt; negative values) on the treadmill belt. Ataxia Coefficient is a within-animal measure of stride-to-stride variability in paw placement (step length), which quantifies gait irregularity. Axis distance and midline distance reflect mediolateral paw placement relative to the animal's direction-of-travel axis and midline, respectively (cm), while overlap distance reflects the relative fore–hind paw placement along the direction of progression. Each dot represents an individual mouse. Data were analyzed by unpaired two-tailed *t*-tests and are presented as mean ± SD.

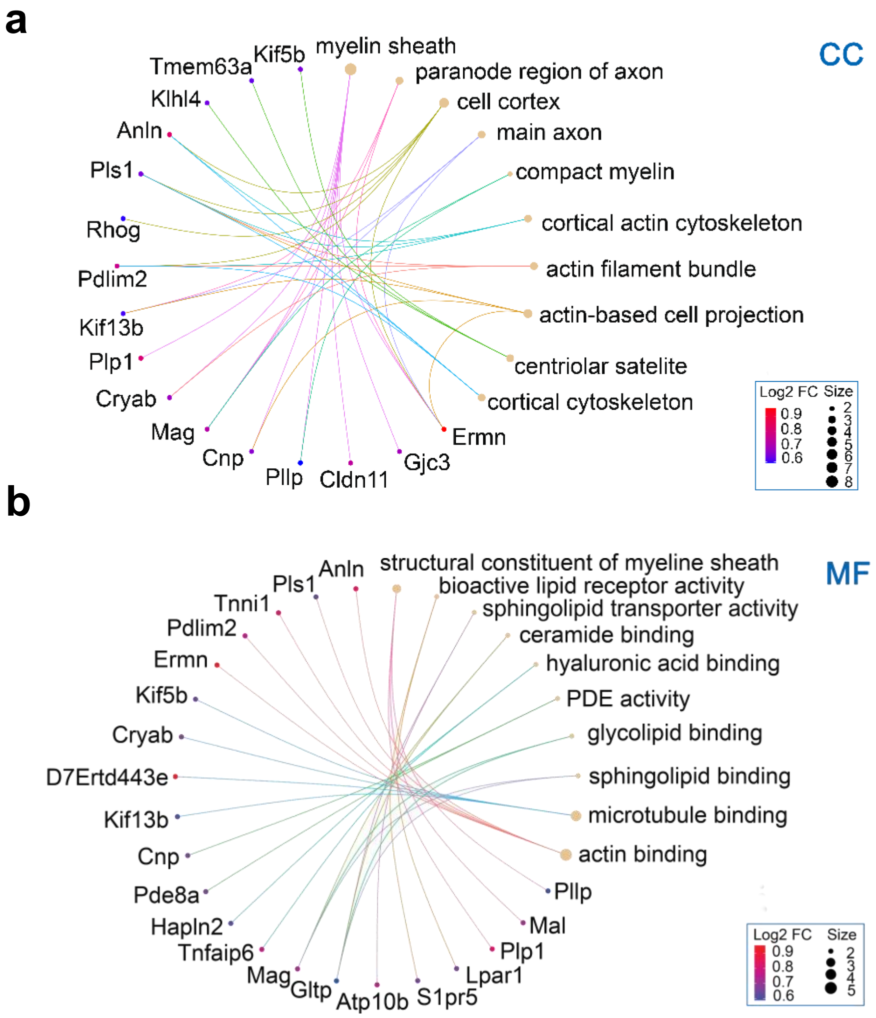


**Supplementary Figure 6. Chord diagrams illustrating the association between differentially expressed genes and enriched Gene Ontology (GO) terms.** Gene-concept networks (cnet) link specific genes of interest to functional categories. Lines connect genes (left arc of each circle) to their annotated GO terms (right arc of each circle). **a** Associations with Cellular Component (CC) terms, highlighting genes related to structures like the myelin sheath, axon, and cytoskeleton. **b** Associations with Molecular Function (MF) terms, highlighting genes involved in activities such as binding (lipid, actin, microtubule) and molecular activities. In both panels, the colored dots next to the gene symbols represent differential expression data. The color gradient indicates the $Log_2$ Fold Change ($Log_2FC$) and the node size reflects gene count per term.
